# Unpredictable chronic stress redistributes novelty-evoked exploratory space use and yields method-dependent identification of resilient-like archetypes in zebrafish

**DOI:** 10.64898/2026.10.29.762878

**Authors:** F. Fuentealba-Villarroel, L.M. Bastos, M. Gallas-Lopes, A.P. Herrmann, D. Zancan, A. Piato

**Affiliations:** Laboratório de Psicofarmacologia e Comportamento (LAPCOM), Departamento de Farmacologia, Instituto de Ciências Básicas da Saúde (ICBS), Universidade Federal do Rio Grande do Sul (UFRGS), Porto Alegre, RS, Brazil; Laboratório de Neurobiologia e Psicofarmacologia Experimental (PsychoLab), Departamento de Farmacologia, Instituto de Ciências Básicas da Saúde (ICBS), Universidade Federal do Rio Grande do Sul (UFRGS), Porto Alegre, RS, Brazil; Laboratório de Neurobiologia Comparada, Departamento de Fisiologia, Instituto de Ciências Básicas da Saúde (ICBS), Universidade Federal do Rio Grande do Sul (UFRGS), Porto Alegre, RS, Brazil; Meta-Research Innovation Center at Stanford (METRICS), Stanford University, Stanford, CA, USA; Programa de Pós-Graduação em Neurociências, Instituto de Ciências Básicas da Saúde (ICBS), Universidade Federal do Rio Grande do Sul (UFRGS), Porto Alegre, RS, Brazil; Programa de Pós-Graduação em Farmacologia e Terapêutica, Instituto de Ciências Básicas da Saúde (ICBS), Universidade Federal do Rio Grande do Sul (UFRGS), Porto Alegre, RS, Brazil

**Keywords:** Unpredictable chronic stress, zebrafish, behavioural resilience, novel tank test, individual variability, behavioural phenotyping

## Abstract

Unpredictable chronic stress (UCS) produces heterogeneous behavioural outcomes, but identifying resilient individuals remains challenging because post-stress behaviour is multidimensional and classification depends on the analytical framework. We characterised novelty-evoked exploration in 178 adult zebrafish (89 unstressed reference, 89 UCS-exposed) using four Novel Tank Test variables: distance travelled, line crossings, upper-zone entries and upper-zone time, from a historical LAPCOM dataset from four independent cohorts. Behaviour was robustly normalised to control distributions within cohorts. In the unstressed population, principal component analysis revealed a dominant exploratory gradient (PC1 = 74.9%; PC2 = 18.7%), whereas weighted *k*-nearest-neighbour Louvain analysis yielded a stable two-profile partition. However, null-model inference and resampling did not support discrete behavioural populations, indicating a predominantly continuous reference architecture. UCS significantly redistributed individuals within this architecture (PERMANOVA: *F*(1,173) = 8.10, *R²* = 0.0423, *p* = 0.0016), reducing occupancy of the higher-exploration reference profile from 55.1% to 24.7%, while multivariate dispersion, Mahalanobis distance and global correlation structure showed no detectable differences. *De novo* Louvain analysis identified lower- and higher-exploration configurations, but correspondence with the control reference was asymmetric: all 44 lower-exploration fish mapped to the lower reference profile, whereas only 22 of 45 higher-exploration fish mapped to the higher reference profile. Thus, higher exploration after UCS was insufficient to define a resilient-like response. Across five analytical frameworks, resilient assignments ranged from 9 to 45 individuals and susceptible assignments from 23 to 68, with partial individual-level convergence. These findings support reference-integrated behavioural stratification for identifying resilient- and susceptible-like candidates for physiological, cellular and molecular validation.

**Highlights:**

I. Unstressed zebrafish showed a continuous multivariate behavioural architecture.
II. UCS redistributed exploration without creating a distinct behavioural space.
III. Higher post-UCS exploration alone did not identify resilient-like zebrafish.
IV. Resilience-like classification depended substantially on the analytical method.
V. Control-referenced stratification identified candidates for mechanistic studies.

## 1. Introduction

Chronic stress does not produce uniform behavioural outcomes. Even under comparable experimental conditions, exposed individuals may differ substantially in the magnitude, organisation and persistence of their responses, creating a central challenge for the experimental study of stress susceptibility and resilience. This heterogeneity is particularly important when behavioural phenotypes are subsequently used to investigate neuroendocrine, cellular or molecular mechanisms, because mechanistic comparisons depend on how resilient- and susceptible-like individuals are initially identified. In zebrafish (*Danio rerio*), unpredictable chronic stress (UCS) provides a well-established experimental framework in which repeated heterogeneous stressors produce behavioural, endocrine and molecular alterations, including changes in exploratory-related behaviour, memory, neuroendocrine signalling and cortisol (Gallas-Lopes et al., 2026, 2023; Piato et al., 2011). However, population-level effects do not establish that all exposed individuals respond similarly, nor do they specify how heterogeneous post-UCS behavioural outcomes should be operationally stratified.

Zebrafish naturally inhabit shallow floodplain environments, including ponds, ditches and water bodies associated with rice cultivation, where they occur in both open water and vegetated margins and occupy the full water column (Spence et al., 2008, 2006). These observations provide an ecological basis for considering spatial deployment as an intrinsic component of zebrafish behaviour, while offering little support for interpreting bottom occupancy as a constitutive habitat preference. In laboratory novelty paradigms, however, the animal does not voluntarily approach an unfamiliar environment but is transferred directly into it. The novel tank test (NTT) therefore imposes spatial novelty and elicits a progressive redistribution of behaviour within an unfamiliar three-dimensional environment. To distinguish this directly observable phenomenon from broader constructs such as anxiety-like behaviour, boldness or curiosity, here we use the term *novelty-evoked exploratory space use* to describe the coordinated deployment of locomotion, spatial occupancy and transitions during exposure to an unfamiliar environment.

Novelty-evoked exploratory space use is neither a unitary measure nor a temporally static behavioural state. During continued exposure to a novel tank, adult zebrafish exhibit time-dependent changes in upper-zone occupancy and in the number of transitions into the upper zone, whereas other behavioural components, including freezing and erratic movement, follow partially dissociable trajectories (Wong et al., 2010). Pharmacological manipulations and exposure to aversive stimuli, such as alarm substance, further show that spatial occupancy, transitions and locomotor or defensive responses need not change in parallel, indicating that these measures cannot be treated as interchangeable readouts of a single underlying construct (Wong et al., 2010). Spatial organisation also extends beyond vertical position. In novelty-based arenas, zebrafish can establish localised areas that are disproportionately occupied and repeatedly revisited, a pattern described as homebase behaviour (Rosemberg et al., 2011; Stewart et al., 2010). These observations distinguish related but non-equivalent properties of behaviour under novelty, including spatial sampling and redistribution, recurrent organisation of space use, and time-dependent behavioural reorganisation during continued exposure.

Furthermore, tank geometry, illumination, wall properties and other characteristics of the testing environment can modify the resulting spatial phenotype, as discussed in greater detail in the review by Golushko et al. (Golushko et al., 2025). Thus, behavioral responses in the NTT are better understood as dynamic interactions between the animal, elapsed experience and the sensory-spatial properties of the testing environment, rather than as static behavioural endpoints.

The neural organisation underlying exploratory and novelty-related behaviour is apparent across developmental stages and becomes increasingly evident at the level of forebrain recruitment in adults. In larval zebrafish, apparently spontaneous locomotion is already temporally structured: successive turns are organised into directional sequences rather than independent random events, and perturbation of the anterior rhombencephalic turning region alters both turning bias and the dependence of current actions on previous actions (Dunn et al., 2016). Although this paradigm differs from the adult NTT, it demonstrates that even early exploratory locomotion contains intrinsic neural organisation extending beyond movement quantity alone. In adult zebrafish, Trotha et al., showed that acute amphetamine recruits neurons in the medial (Dm) and lateral (Dl) pallial territories, whereas Dm is also activated during drug-seeking approach and light-avoidance behaviour, providing evidence that behaviours involving motivational and valence-related processing engage defined pallial circuits (Trotha et al., 2014). This adult forebrain involvement is directly extended to novelty exposure by Rajput et al., who mapped whole-brain *c-fos* activity following the NTT and identified increased activity across multiple telencephalic regions, including pallial and subpallial territories, within a distributed functional network spanning preoptic, thalamic, hypothalamic, olfactory and sensorimotor regions (Rajput et al., 2025). Network analysis highlighted several ventral telencephalic territories—particularly Vv, Vd-dd, Vd-vd and Vdd—as highly central, while the anterior part of the parvocellular preoptic nucleus (PPa), the olfactory bulb and the mesencephalic nucleus of the trigeminal nerve (Vmn) also emerged as prominent nodes linking major network components (Rajput et al., 2025).

These territories are distinguishable within the established neuroanatomical organisation of the adult zebrafish brain (Kenney et al., 2021; Wullimann et al., 1996), although anatomical correspondence does not by itself establish functional equivalence with mammalian structures. Collectively, these findings are consistent with novelty-evoked exploratory space use emerging from distributed integration of sensory information, internal state, action selection and motor implementation rather than from a single dedicated exploration circuit. This supports treating the NTT not as a single locomotor or anxiety-related readout, but as a biologically organised multidimensional behavioural domain whose internal structure can be used for initial phenotypic stratification.

This multidimensional organisation is accompanied by substantial inter-individual variability. Population and strain differences in responses to novel stimuli have been observed even under common laboratory rearing conditions, indicating that such variability cannot be attributed solely to immediate environmental experience (Moretz et al., 2007; Rajput et al., 2022; Spence et al., 2008; Wright et al., 2003). More directly, three-dimensional analysis of more than 400 adult zebrafish demonstrated that distance travelled, spatial coverage and position within a novel tank combine into heterogeneous behavioural configurations that are not adequately represented by a simple bold–shy dichotomy (Rajput et al., 2022). These configurations were influenced by strain and sex and showed both persistence and reorganisation across repeated exposures, indicating that individual exploratory phenotypes are structured but behaviourally plastic rather than fixed binary types. Individual variation in novelty-evoked exploratory space use may therefore reflect structured differences within a multidimensional behavioural architecture rather than measurement noise or variation along a single exploratory axis.

This property becomes particularly relevant after UCS. A systematic review and meta-analysis of the first decade of zebrafish UCS research found overall increases in anxiety/fear-related measures and cortisol and reduction in locomotor function, whereas no significant summary effect was detected for social behaviour (Gallas-Lopes et al., 2023), a finding later corroborated by Neves et al. (Neves et al., 2026). However, the social-behaviour literature included heterogeneous and sometimes opposing effects across studies, and the pooled null effect should therefore be interpreted cautiously (Gallas-Lopes et al., 2023). A standardised UCS protocol for adult zebrafish employs a 14-day regimen of varied stressors, with an emphasis on reproducibility and control of experimental sources of variation (Gallas-Lopes et al., 2026).

Against this background, the multidimensional, temporally organised, neurobiologically distributed and individually variable nature of novelty-evoked exploratory space use provides a rationale for asking whether the NTT contains sufficient behavioural information to resolve heterogeneous organism-level responses following UCS. Such information is unlikely to reside exclusively in a single endpoint, such as upper-zone time or total distance travelled, because these variables capture different and only partially overlapping dimensions of space use. Their joint organisation may therefore preserve behavioural information that is lost when post-UCS responses are inferred from any single NTT measure considered in isolation.

This does not imply that novelty-evoked exploratory space use exhausts the biological construct of resilience, nor that memory, social behaviour, endocrine responses or other domains are irrelevant to its broader validation. Rather, it establishes the NTT as a defined organism-level behavioural domain in which divergent post-UCS responses can be operationally stratified before their physiological or neurobiological correlates are examined. The principal difficulty then becomes how such responses should be operationalised. Biological resilience is intrinsically dynamic and operates across molecular, cellular, tissue and whole-organism scales through integrated responses that permit biological systems to recover from perturbation or establish adapted states following challenge (Weavers, 2024). At the behavioural level, existing approaches retain markedly different amounts of information and apply different criteria for successful adaptation. Swaminathan et al. classified zebrafish larvae according to the dynamics of behavioural rebound after a major stressor, distinguishing fast and slow recoverers and subsequently demonstrating stability across additional challenges and developmental stages (Swaminathan et al., 2023). Mueller et al., by contrast, identified resilient-like outcomes through multivariate behavioural stratification and interpreted exposed individuals with control-comparable behavioural profiles as resilient (Mueller et al., 2021). These approaches illustrate that resilience is not directly observed as a categorical behavioural trait but inferred from an explicit relationship between exposure, behavioural outcome and a reference criterion. Moreover, in cross-sectional post-UCS designs, a control-like behavioural configuration cannot by itself distinguish resistance to perturbation from recovery after a previous deviation. For a more in-depth conceptual discussion of this distinction, see the Opinion article by Bhatnagar (Bhatnagar, 2021). We therefore use the terms *resilient- and susceptible-like archetypes* to denote operationally defined behavioural responses following UCS rather than independently validated biological phenotypes.

Here, we retrospectively analysed a historical dataset from four independent experimental cohorts, equally distributed between unstressed control and UCS-exposed populations. We first reconstructed a control-derived behavioural reference architecture from four NTT dimensions and used this fixed reference to quantify population-level behavioural redistribution following UCS. We then characterised the internal behavioural architecture of the UCS-exposed population *de novo* using Louvain community detection (Blondel et al., 2008; Rajput et al., 2022). Rather than defining resilient- and susceptible-like archetypes from a single behavioural threshold, the post-UCS communities were initially treated as algorithmic descriptors and then interpreted relative to the independently derived control reference. Finally, we compared this reference-integrated strategy with four alternative operational frameworks—Swaminathan-like thresholding, PCA-derived index quantiles, Gaussian mixture modelling and Two-Step clustering—and quantified their individual-level convergence. This design was intended to determine how UCS redistributes multidimensional novelty-evoked exploratory space use, how inter-individual post-stress heterogeneity relates to the unstressed behavioural reference, and how strongly the operational identification of resilient- and susceptible-like archetypes depends on the analytical framework used. By separating behavioural stratification from subsequent biological validation, this framework provides a phenotypic basis for future investigation of the neuroendocrine, cellular and molecular mechanisms associated with differential adaptation to chronic stress.

## 2. Materials and methods

### 2.1 Subjects and experimental design

This study comprised a retrospective *in silico* analysis of a historical behavioural dataset generated at the Laboratory of Psychopharmacology and Behaviour (LAPCOM). The dataset included 178 adult zebrafish, equally distributed between unstressed control (*n* = 89) and UCS-exposed (*n* = 89) groups. These animals originated from four independent experimental cohorts (Bertelli et al., 2021; Marcon et al., 2019; Neves et al., 2026; Sachett et al., 2022). Cohort-level control/UCS sample sizes were 24/24, 11/12, 18/17 and 36/36 zebrafish, respectively. Individual identity and cohort provenance were retained throughout the analytical workflow.

The historical dataset was used to reconstruct a control-derived reference architecture of novelty-evoked exploratory space use, quantify population-level behavioural redistribution following UCS, characterise *de novo* post-UCS configurations within this multidimensional behavioural space, and operationally identify resilient- and susceptible-like archetypes based on their relationship to the unstressed behavioural reference. The retrospective analysis formed part of a broader research programme documented and made publicly available through the Open Science Framework (OSF; Fuentealba-Villarroel et al., 2025). This public record provides a time-stamped account of the overarching scientific rationale and analytical framework of the research programme.

Four variables measured in the novel tank test defined the multidimensional novelty-evoked exploratory space use: total distance travelled, number of line crossings between the virtual horizontal zones, number of entries into the upper zone, and time spent in the upper zone. These variables were used to reconstruct the control-derived behavioural reference architecture and to characterise UCS-associated changes in population organisation.

### 2.2 Behavioural data preprocessing and control-referenced normalisation

Behavioural data from the historical dataset were processed while preserving the original structure of the four experimental cohorts and individual traceability. Because numerical animal identifiers were not unique across cohorts, each zebrafish was uniquely identified using a composite identifier combining its cohort of origin and the original numerical identifier. This identity was retained throughout data preprocessing and downstream analyses.

To place behavioural measurements obtained across independent experimental cohorts onto a common analytical scale while retaining the biological reference provided by unstressed animals, each behavioural variable was robustly normalised relative to its cohort-specific control distribution. For each variable and within each cohort, the median and raw median absolute deviation (MAD) were estimated exclusively from control zebrafish. Values from both control and UCS-exposed zebrafish belonging to the corresponding cohort were then transformed as:

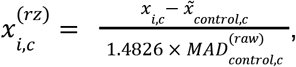

where *i* denotes the individual zebrafish, *c* denotes the experimental cohort, *x_i_*_,*c*_ represents the observed behavioural value for each individual, *~x_control,c_* represents the median calculated from control zebrafish within the corresponding cohort, 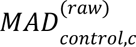 represents the raw median absolute deviation of the same control distribution and 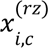 represents the resulting control-referenced robust *z*-score. Unlike a conventional *z*-score, this transformation uses the control median and the scaled median absolute deviation rather than the arithmetic mean and standard deviation. The factor 1.4826 scales the raw MAD to provide a robust estimate of the standard deviation under an approximately Gaussian distribution. Importantly, UCS-exposed zebrafish did not contribute to the estimation of either the reference centre or scale. The same control-derived parameters were applied unchanged to control and UCS-exposed zebrafish from the corresponding cohort.

This procedure was independently applied to the four behavioural variables defining the historical multidimensional novelty-evoked exploratory space use. The resulting control-referenced robust coordinates were used throughout the reconstruction of the reference behavioural architecture, projection of UCS-exposed zebrafish and *de novo* characterisation of behavioural structure within the UCS-exposed population. No subsequent standardisation based on the distribution of the UCS-exposed population was performed.

### 2.3 Reconstruction of the behavioural reference architecture

#### 2.3.1 Control-derived reference architecture of novelty-evoked exploratory space use

The behavioural reference architecture was reconstructed exclusively from the 89 control zebrafish using the four cohort-specific, control-referenced robust coordinates described above. Following normalisation, zebrafish from the four independent cohorts were analysed jointly, while cohort and individual identity were retained throughout the analytical workflow. No additional standardisation was applied at this stage.

Principal component analysis (PCA) was first used to characterise the dominant multivariate axes of behavioural variation within the control population. PCA was performed on the complete four-dimensional robust behavioural matrix after arithmetic mean-centring, without additional variance scaling. The resulting component loadings, scores and explained variance defined a continuous projection-ready representation of the control behavioural space. PCA was used to describe multivariate organisation and dimensional structure, rather than to impose discrete behavioural categories.

To investigate whether local organisation was detectable within this reference space, a weighted *k*-nearest-neighbour (*k*NN) graph was constructed directly from the complete four-dimensional robust coordinates (Blondel et al., 2008; Rajput et al., 2022). Pairwise dissimilarity was quantified using Euclidean distance, and edge similarity was defined as:

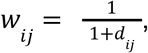

where *d_ij_* denotes the Euclidean distance between zebrafish *i* and *j*. The resulting graph was defined as the weighted, undirected union of individual *k*-nearest-neighbour relations. Candidate neighbourhood sizes were systematically evaluated over *k* = 5–80. For each *k*, Louvain community detection was repeated across 100 reproducible random seeds to evaluate sensitivity to stochastic optimisation. A representative partition was selected from the repeated solutions according to its Adjusted Rand Index (ARI) similarity to the remaining partitions, with modularity and deterministic seed ordering used only as secondary tie-breakers. Candidate graph solutions were further characterised using mean silhouette width, the Calinski–Harabasz index, the Davies–Bouldin index and weighted modularity.

Because graph-partitioning algorithms can identify apparent structure even in continuous multivariate distributions, evidence for discrete organisation was evaluated independently using formal null-model inference. The primary null model was constructed in data space using a Gaussian-copula procedure that exactly preserved the empirical marginal distribution of each robust behavioural variable while approximately retaining their rank-dependence structure. For each of 1,000 null datasets, the complete *k* = 5–80 optimisation and partition-selection procedure was repeated, thereby incorporating hyperparameter selection into the null distribution of weighted modularity. Complementary analyses included fixed-*k* data-space comparisons, a family-wise maximum-standardised-modularity assessment across the evaluated *k* range and degree-preserving graph rewiring. Thus, algorithmic segmentation and evidence for genuine behavioural discreteness were explicitly treated as distinct inferential questions.

The exact-equivalence plateau extended from *k* = 44 to *k* = 46. The central value, *k* = 45, was retained as the representative configuration for null-model and resampling analyses. For subsequent projection analyses, the identical two-segment partition obtained at *k* = 46 was retained as the authorised fixed configuration. Because all values within this plateau produced identical individual-level memberships, the use of *k* = 46 did not represent a distinct solution. The two algorithmic segments were subsequently oriented according to their multivariate exploratory profiles and operationally designated as higher- and lower-exploration reference profiles. These designations described positions within the control-derived behavioural continuum and were not interpreted as independently validated biological subtypes.

The conditional reproducibility of the retained graph partition was additionally assessed by case-bootstrap resampling. Control zebrafish were sampled with replacement while their previously established robust coordinates remained fixed, and the weighted union-*k*NN graph was reconstructed using *k* = 45, the representative configuration retained for resampling analyses. Agreement between each bootstrap partition and the retained full-sample partition was quantified primarily by the ARI calculated across the unique original zebrafish represented in each resample. When an individual occurred more than once, its bootstrap membership was determined by modal assignment across resampled copies, with deterministic resolution of ties. An ARI calculated across all bootstrap rows was retained as a secondary diagnostic. This procedure assessed the algorithmic and sampling stability of the retained segmentation and did not involve re-optimisation of *k*.

#### 2.3.2 Projection of UCS-exposed zebrafish onto the control-derived reference architecture

Population-level behavioural multidimensional novelty-evoked exploratory space use reorganisation following UCS was evaluated by projecting the 89 UCS-exposed zebrafish into the fixed control-derived behavioural space. No PCA, normalisation parameters or Louvain partition was re-estimated from the UCS population for this analysis. Instead, each UCS zebrafish retained the four cohort-specific control-referenced robust coordinates established during preprocessing and was projected onto the PCA axes defined exclusively by the control population using the fixed control PCA centre and rotation matrix.

The operational control-derived reference profiles identified above were held fixed. Control zebrafish retained their original Louvain-derived reference-profile memberships, whereas each UCS-exposed zebrafish was assigned to the nearest fixed control-derived reference-profile centroid according to Euclidean distance in the complete four-dimensional robust behavioural space. The fidelity of this centroid-based projection rule within the reference population was assessed using leave-one-out control centroids, thereby preventing each control zebrafish from being assigned relative to a centroid to which it had contributed. Importantly, no Louvain community detection was performed on the UCS-exposed population at this stage; all UCS assignments therefore represented projections onto the pre-existing control-derived reference architecture.

Complementary multivariate analyses were used to quantify the displacement of the UCS-exposed population relative to the control-derived reference. Overall four-dimensional centroid displacement (*D*₄D) was calculated as the Euclidean distance between the control and UCS centroids in the complete four-dimensional robust behavioural space. A treatment-associated difference in multivariate location was evaluated using partial permutational multivariate analysis of variance (PERMANOVA; Anderson, 2001; Anderson, 2017), with cohort included as a conditioning factor and 9,999 permutations restricted within cohort. Because PERMANOVA can be influenced by differences in within-group dispersion, multivariate dispersion was evaluated separately using PERMDISP, based on distances to the group spatial median and 9,999 residual permutations restricted within cohort (Anderson, 2006).

Departure from the control reference was further quantified using Mahalanobis distance in the pooled control-referenced space. The control reference centre and classical covariance matrix were estimated from the pooled control population after cohort-specific normalisation. For control zebrafish, distances were obtained using leave-one-out reference estimates, whereas UCS distances were calculated relative to the complete control reference. Square-root Mahalanobis distances were used for interpretation. The primary group comparison reconstructed the complete reference-estimation procedure under label permutation, thereby incorporating uncertainty associated with construction of the control reference into the null distribution.

Finally, potential reorganisation of inter-variable relationships was examined independently of the graph segmentation. Spearman correlation matrices were calculated separately for the complete control and UCS populations. The global difference between correlation structures was quantified across the six unique off-diagonal pairwise correlations and assessed by permutation of pooled, within-group standardised midrank scores. Pairwise changes in Spearman correlation were retained as secondary analyses, with maximum-statistic adjustment used to control the family-wise error rate across the six associations.

#### 2.3.3 De novo organisation of novelty-evoked exploratory space use within the UCS-exposed population

The internal organisation of novelty-evoked exploratory space use within the UCS-exposed population was subsequently investigated *de novo* using only the 89 UCS-exposed zebrafish. The cohort-specific, control-referenced robust coordinates established during preprocessing were retained unchanged, and no UCS-specific normalisation or rescaling was performed. This restriction applied to the behavioural coordinate system itself and did not preclude the arithmetic mean-centring used solely to fit the descriptive PCA. Accordingly, control zebrafish contributed only the reference coordinate system in which UCS behaviour was expressed; neither their higher- and lower-exploration reference-profile memberships nor their cluster structure were imposed on the *de novo* UCS analysis.

A separate PCA was first fitted to the four robust coordinates of the UCS population to describe its dominant internal axes of behavioural variation. Variables were arithmetic mean-centred within the UCS population but were not additionally scaled. This centring defined the origin of the descriptive PCA without altering the inherited robust units or pairwise Euclidean geometry. As in the control analysis, PCA served solely as a representation of internal multivariate variation and was not itself used to define behavioural communities.

*De novo* community detection was then performed directly in the complete four-dimensional inherited robust space. Euclidean distances were converted to similarities as:

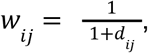

and weighted, undirected union-*k*NN graphs were constructed over *k* = 5–80. Louvain community detection was applied to each candidate graph using a reproducible random seed (Blondel et al., 2008; Rajput et al., 2022). Candidate solutions were evaluated for both internal separation and stability across neighbouring values of *k*. A reproducible candidate region was defined as the longest consecutive *k* plateau retaining the same number of communities and showing high agreement between adjacent partitions, quantified by ARI (Rajput et al., 2022). An ARI threshold of 0.95 was applied primarily and 0.90 secondarily if necessary. Within the selected plateau, the representative *k* corresponded to the medoid partition, defined by maximal mean ARI to the remaining plateau partitions; distance to the plateau midpoint, silhouette width and the smaller *k* were used sequentially as deterministic tie-breakers.

The stability of the resulting candidate partition was subsequently evaluated independently from *k* selection. Algorithmic stability was assessed by repeated Louvain optimisation on the fixed selected graph, whereas sampling stability was examined through case-bootstrap resampling with replacement and reconstruction of the same graph topology rule at the selected *k*.

Bootstrap agreement with the original partition was quantified primarily using ARI across unique original zebrafish, with repeated observations of the same individual collapsed by modal community assignment; row-level ARI was retained as a secondary diagnostic. The selected *k* was not re-optimised during these stability analyses.

At this stage, community labels were treated as algorithmic descriptors of the internal organisation of the UCS population. Numerical Louvain labels were reordered according to mean UCS PC1 solely to provide deterministic labelling and carried no a priori biological meaning. Resilience-like and susceptibility-like archetype interpretations were assigned only subsequently, by comparing the *de novo* UCS communities with the fixed higher- and lower-exploration reference profiles derived from the control population. These interpretations were operational and were not considered evidence of independently validated biological phenotypes.

### 2.4 Analytical frameworks for classifying resilience-like and susceptibility-like archetypes

Resilience-like and susceptibility-like behavioural responses to UCS were operationalised using five analytical classification frameworks that interrogated the multidimensional behavioural archetype from complementary statistical perspectives. Classification was based exclusively on behavioural information. Control zebrafish provided the behavioural reference against which UCS-exposed individuals were evaluated, whereas archetype classification was performed within the UCS-exposed population.

#### 2.4.1 Behavioural Index

A continuous Behavioural Index (BI) was constructed to quantify the position of each UCS-exposed zebrafish along the multivariate axis linking the UCS and control behavioural centroids. PCA was fitted exclusively to the control-referenced behavioural coordinates of control zebrafish, and both control and UCS-exposed zebrafish were subsequently represented in this fixed PCA space. All available principal components were retained, thereby preserving the complete multidimensional information contained in the behavioural dataset rather than restricting the index to the first two components used for graphical representation.

Within this PCA space, the complete PCA-score vector for zebrafish *i* was defined as:

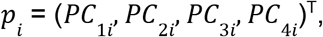

where T denotes vector transposition. The direction from the UCS centroid towards the control centroid was then used to construct the BI. For each principal component *j*, the centroid difference was calculated as:

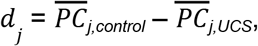

These differences were normalised to obtain a unit-length directional vector:

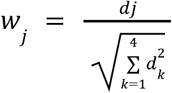

The BI of each zebrafish was subsequently calculated as the scalar projection of its PCA-score vector onto this control-directed axis:

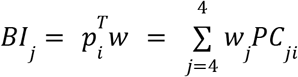

The resulting weights were −0.976833 for PC1, −0.069327 for PC2, −0.190533 for PC3 and 0.068468 for PC4. Accordingly, the implemented index was:

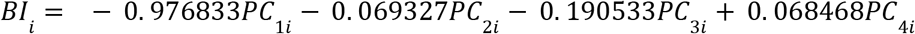

The weights, rather than the resulting BI values, were normalised. Higher BI values indicated positions further along the control-directed behavioural axis, whereas lower values indicated positions further along the UCS-directed axis. The BI was therefore interpreted as a continuous directional score rather than as a measure of Euclidean proximity to the control population, an independent biological marker or an intrinsically categorical archetype.

The BI provided a common and interpretable axis for comparing the classification frameworks. It was used directly to construct the Swaminathan-like and PCA-quantile classifications and was used only after model fitting to orient the classes obtained through Gaussian mixture modelling, Louvain community detection and Two-Step clustering. The BI was not included as an additional input variable in these clustering procedures because it was mathematically derived from the same multidimensional behavioural information.

#### 2.4.2 Swaminathan-like classification

The first analytical framework was based on an adaptation of the behavioural-resilience approach proposed by Swaminathan et al., 2023. UCS-exposed zebrafish were classified according to their position along the BI relative to a control-referenced behavioural threshold. The threshold was defined by jointly optimising behavioural separation, quantified by Cohen’s *d*, and control–UCS discrimination, quantified by Fisher’s exact test; both metrics were min–max normalised to a 0–1 scale and summed, with the threshold yielding the highest combined score selected for classification (for methodological details see Swaminathan et al., 2023). Individuals with BI values above the selected threshold were classified as resilient-like archetypes, whereas those with values below the threshold were classified as susceptible-like archetypes.

This framework generated a binary classification and therefore did not impose an intermediate category. Its inclusion was intended to represent a threshold-based operational definition of resilience in which preservation of a control-like behavioural archetype following UCS constitutes the principal criterion.

#### 2.4.3 PCA-quantile classification

A second framework was constructed directly from the distribution of the BI within the UCS-exposed population. UCS zebrafish were ranked according to their BI values and classified using the first and third quartiles of the UCS distribution as empirical boundaries. Individuals in the lower quartile were classified as susceptible-like archetype, those in the upper quartile as resilient-like archetype, and zebrafish located within the interquartile range as intermediate behaviour.

This procedure generated a deliberately distribution-based classification without assuming an underlying parametric mixture or discrete cluster structure. By definition, it identified the two behavioural extremes of the UCS population while retaining the central 50% as an intermediate archetype (Espigares et al., 2025; Leong et al., 2025; Yata and Aoshima, 2020).

#### 2.4.4 Gaussian mixture modelling

Model-based classification was performed using Gaussian mixture modelling (GMM; Fraley and Raftery, 2007; Videlier et al., 2015). The model was fitted to the complete set of PCA scores representing the control-referenced behavioural coordinates rather than to the BI itself. The BI was explicitly excluded from model fitting because it was mathematically derived from the same multidimensional behavioural information and its inclusion would therefore have introduced redundant information.

Candidate Gaussian mixture models differing in covariance parameterisation and number of components were compared using the Bayesian Information Criterion (BIC). The range of candidate components was sufficiently broad to avoid imposing a three-component solution a priori. Model uncertainty was additionally evaluated using the Integrated Completed Likelihood (ICL) and posterior classification probabilities.

The selected GMM identified three latent Gaussian components. Following model fitting, these components were ordered according to their mean BI along the common control-referenced behavioural continuum. The component with the lowest mean BI was operationally designated as susceptible-like, the component with an intermediate mean BI as intermediate, and the component with the highest mean BI as resilient-like. These designations were applied solely to orient the independently derived latent components relative to the control-referenced direction of novelty-evoked exploratory space use. The BI did not contribute to model estimation, component selection or individual component assignment.

#### 2.4.5 Louvain community detection

A graph-based classification was performed using Louvain community detection (Beigloo et al., 2024; Blondel et al., 2008; Rajput et al., 2022). Unlike the GMM procedure, Louvain analysis was conducted directly in the complete control-referenced behavioural coordinate space.

Pairwise Euclidean distances between UCS-exposed zebrafish were converted to similarities according to:

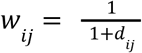

Weighted, undirected union-*k*-nearest-neighbour graphs were then constructed, and Louvain community detection was applied across candidate neighbourhood sizes. Selection of the representative graph configuration was based on the stability of neighbouring partitions rather than on a single isolated optimum. Candidate regions were identified from consecutive *k* values producing stable community structure, with agreement between neighbouring partitions quantified using the ARI. A representative *k* was subsequently selected within the stable region according to the partition most representative of the surrounding solutions, using predefined deterministic tie-breaking criteria when required.

The resulting Louvain communities were algorithmic groups and were not assumed a priori to represent resilient-, intermediate- or susceptible-archetypes. For the cross-method comparison, the communities were ordered according to their mean BI and operationally labelled resilient-like or susceptible-like archetype. The BI was used only for *post hoc* orientation and did not contribute to graph construction or community detection. These method-specific labels were distinct from the stricter reference-integrated classification described above.

#### 2.4.6 Two-Step cluster analysis

A complementary Two-Step cluster analysis was performed in IBM SPSS Statistics (version 27.0.1) using the complete set of control-referenced behavioural coordinates from the UCS-exposed population (Benassi et al., 2020; IBM Corp., 2025; Mueller et al., 2021).

Log-likelihood-based distance was used, and the number of clusters was selected automatically according to the Bayesian Information Criterion (BIC), with a predefined maximum of 15 candidate clusters. No additional standardisation was applied because all variables had already been transformed to the common control-referenced robust scale described above.

The resulting two clusters were ordered according to their mean BI. The cluster with the lower mean BI was operationally classified as susceptible-like archetype, whereas the cluster with the higher mean BI was operationally classified as resilient-like archetype, according to Mueller et al. (Mueller et al., 2021). The BI was used only for *post hoc* orientation and did not contribute to the Two-Step clustering procedure.

#### 2.4.7 Cross-method agreement and behavioural consensus

Because the five analytical classification frameworks differed in their statistical assumptions and operational definitions of resilience, agreement among methods was evaluated rather than assuming that any single framework constituted a definitive ground truth. Pairwise agreement was quantified using the proportion of exact agreement, Cohen’s *κ* and the ARI. Association between classifications was quantified using Cramér’s *V* and Fisher’s exact test. Monte Carlo estimation was used when sparse cell counts made asymptotic approximations inappropriate.

A behavioural consensus classification was then constructed from the five method-specific class assignments obtained for each UCS-exposed zebrafish. Zebrafish assigned to the same archetype by all five frameworks were designated as unanimous resilient-like or unanimous susceptible-like archetype. Agreement of 4/5 frameworks was considered high consensus for the corresponding archetype. Zebrafish failing to reach this criterion were retained as behaviourally ambiguous/intermediate archetypes rather than being forcibly assigned to one of the two extreme archetypes.

This consensus procedure was deliberately conservative. This procedure was designed to identify zebrafish whose classifications were concordant across analytical frameworks with different statistical assumptions, while explicitly retaining uncertainty for individuals whose classification depended on the analytical framework employed. Cross-method consensus was interpreted as agreement among method-specific operational definitions rather than as independent replication or biological validation.

### 2.5 Statistical analysis and computational implementation

#### 2.5.1 General analytical framework

Statistical analyses were conducted primarily in R (version 4.6.1, R Core Team, 2026), with IBM SPSS Statistics (version 27.0.1) used specifically for the Two-Step cluster analysis. All analyses were performed at the individual-zebrafish level. Statistical tests were two-sided unless otherwise specified, and statistical significance was defined as *p* < 0.05. Exact or permutation-based inference was preferentially used when distributional assumptions, sparse contingency tables or the structure of the analytical procedure made conventional asymptotic inference inappropriate.

Continuous behavioural variables were summarised using means and standard deviations when required for model-based procedures and medians and interquartile ranges when distributional characteristics favoured robust descriptive statistics. Effect sizes were reported together with inferential statistics whenever appropriate. The analytical strategy explicitly distinguished between hypothesis-testing procedures, measures of effect magnitude, algorithmic optimisation and assessments of classification stability; optimisation criteria were therefore not interpreted as inferential evidence of biological group separation. Between-class contrasts involving behavioural variables used to construct or orient the classifications were interpreted exclusively as descriptive characterisations of the resulting partitions and not as independent validation of those classifications.

Reproducibility was maintained by fixing random seeds before stochastic procedures, including permutation tests, bootstrap resampling, Louvain optimisation and Gaussian mixture modelling where applicable. Intermediate analytical objects, individual-level classifications and final statistical outputs were retained to permit reconstruction of the complete analytical workflow.

#### 2.5.2 Statistical inference for the dataset

Statistical inference for population-level differences between control and UCS-exposed zebrafish was conducted in the four-dimensional control-referenced behavioural space.

Treatment-associated differences in multivariate location and dispersion were evaluated using partial PERMANOVA and PERMDISP (Anderson, 2017, 2006, 2001), respectively, following the cohort-restricted permutation procedures described above. These complementary analyses were used to distinguish treatment-associated displacement in novelty-evoked exploratory space use from differences in within-group multivariate dispersion.

Multivariate dispersion was therefore examined independently using distances to the group spatial median in the same four-dimensional behavioural space, with permutation-based inference preserving the original cohort structure. Distance from the control-derived behavioural reference was additionally quantified using Mahalanobis distance. Control distances were obtained using leave-one-out reference estimates, whereas UCS-exposed zebrafish were evaluated relative to the complete control reference. The primary permutation test reconstructed the control reference after treatment-label permutation rather than treating the originally estimated reference parameters as fixed, thereby propagating reference-estimation uncertainty into the null distribution.

Differences in behavioural correlation structure were evaluated using Spearman’s rank correlations coefficient (*r_s_*). Correlation matrices were calculated separately for control and UCS-exposed zebrafish, and differences between corresponding correlations were expressed as:

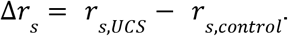

A global permutation statistic incorporating the six unique pairwise correlations among the four behavioural variables was used to test whether the overall correlation architecture differed between populations. Pairwise Δ*r_s_* values were subsequently examined as secondary analyses, with maximum-statistic adjustment used to control the family-wise error rate across the six correlations. Holm-adjusted probabilities were retained as an additional multiplicity-sensitive diagnostic.

For graph-based analyses, modularity, silhouette width, the Calinski–Harabasz index and the Davies–Bouldin index were used as descriptive optimisation criteria and were not interpreted as conventional hypothesis tests. Stability among alternative Louvain solutions was quantified using the ARI. Evidence that the control-derived reference graph segmentation exceeded that expected from a continuous multivariate distribution was evaluated using the formal null-model procedures described above, with 1,000 complete null reconstructions of the optimisation procedure. Bootstrap stability was similarly quantified using ARI relative to the retained full-sample partition.

#### 2.5.3 Comparison among behavioural classification frameworks

Agreement among the five analytical classification frameworks was evaluated at the individual-zebrafish level. Pairwise exact agreement quantified the proportion of zebrafish receiving identical classifications under two methods. Because exact agreement alone does not account for chance agreement or the structure of multicategory assignments, concordance was additionally quantified using Cohen’s κ, ARI and Cramér’s *V*.

Statistical dependence between classifications was assessed using Fisher’s exact test. Monte Carlo estimation was used for contingency tables containing sparse cells for which exact enumeration was computationally impractical or asymptotic χ^2^ approximations were inappropriate. Agreement metrics were interpreted jointly rather than relying on a single statistic, because the five analytical frameworks differed in the number, prevalence and geometry of their resulting classes.

The cross-method consensus classification was derived directly from the five method-specific class assignments. Agreement by at least four of the five frameworks defined high-consensus resilient or susceptible archetypes, whereas five concordant classifications defined unanimous resilient or susceptible archetypes. Zebrafish not fulfilling either high-consensus criterion were retained as ambiguous/intermediate.

#### 2.5.4 Software, coding and computational reproducibility

Data management, robust normalisation, principal component analyses, distance calculations, permutation procedures, bootstrap analyses, graph construction, Louvain community detection, Gaussian mixture modelling, statistical inference and figure preparation were implemented using scripted workflows in R (version 4.6.1, R Core Team, 2026). The Two-Step cluster analysis was performed separately in IBM SPSS Statistics (version 27.0.1) using log-likelihood distance and automatic BIC-based selection of the number of clusters, with the maximum number of clusters set to 15 and without additional standardisation of the previously normalised behavioural variables.

All stochastic analyses used explicitly defined random seeds, and analytical objects were saved at successive stages of the workflow to prevent inadvertent re-estimation of previously validated transformations or classifications. In particular, downstream analyses inherited the previously established control-referenced coordinates rather than recalculating normalisation parameters from transformed or subgrouped datasets. This ensured analytical traceability from raw individual behavioural measurements through reference-space construction and behavioural classification.

Data processing, computational analyses and statistical inference were performed in R version 4.6.1 under Microsoft Windows 10. Data manipulation was conducted principally using dplyr (v1.2.1; Wickham et al., 2026), tidyr (v1.3.2; Wickham et al., 2025) and tibble (v3.3.1; Müller et al., 2026), whereas graphical visualisation was performed using ggplot2 (v4.0.3; Wickham et al., 2026b), ggrepel (v0.9.8; Slowikowski et al., 2026) and patchwork (v1.3.2; Pedersen, 2025). Data import and export were handled using openxlsx (v4.2.8.1; Schauberger et al., 2025).

Graph-based analyses, including construction and analysis of weighted k-nearest-neighbour networks and Louvain community detection, were implemented using igraph (v2.3.3; Csárdi et al., 2026), with cluster (v2.1.8.2; Maechler et al., 2026) used for complementary internal clustering metrics. Gaussian mixture modelling was performed using mclust (v6.1.2; Scrucca et al., 2023). PERMANOVA and multivariate dispersion analyses were implemented using vegan (v2.7.5; Oksanen et al., 2026) and permute (v0.9.10; Simpson et al., 2026), whereas irr (v0.85; Gamer et al., 2026) and mclust were used for complementary agreement metrics among behavioural classification frameworks. Additional statistical procedures and effect-size calculations were implemented using base R functions and rstatix (v1.0.0; Kassambara, 2026).

Analysis scripts, the original behavioural dataset, data dictionary, computational environment records, execution order and SPSS Two-Step materials are publicly archived in Zenodo (https://doi.org/10.5281/zenodo.23425599). The version-controlled project repository is maintained at GitHub (https://github.com/MisfoldERMind/zebrafish-ucs-exploratory-archetypes).

OpenAI ChatGPT was used during manuscript preparation to assist development and review of analysis code, and Anthropic Claude was used to assist adaptation of figure-generation scripts to the journal’s formatting requirements. All analytical decisions, code execution, verification of results, interpretation of findings, and final manuscript content were critically reviewed and approved by the authors, who take full responsibility for the work.

## 3. Results

### 3.1 Novelty-evoked exploratory space use in the unstressed reference population forms a continuum with an operational two-profile partition

To establish the reference behavioural architecture against which behavioural configurations observed after unpredictable chronic stress could subsequently be interpreted, we first examined the organisation of individual behavioural variation in unstressed zebrafish in the NTT (Fig. 1).

**Figure 1.**
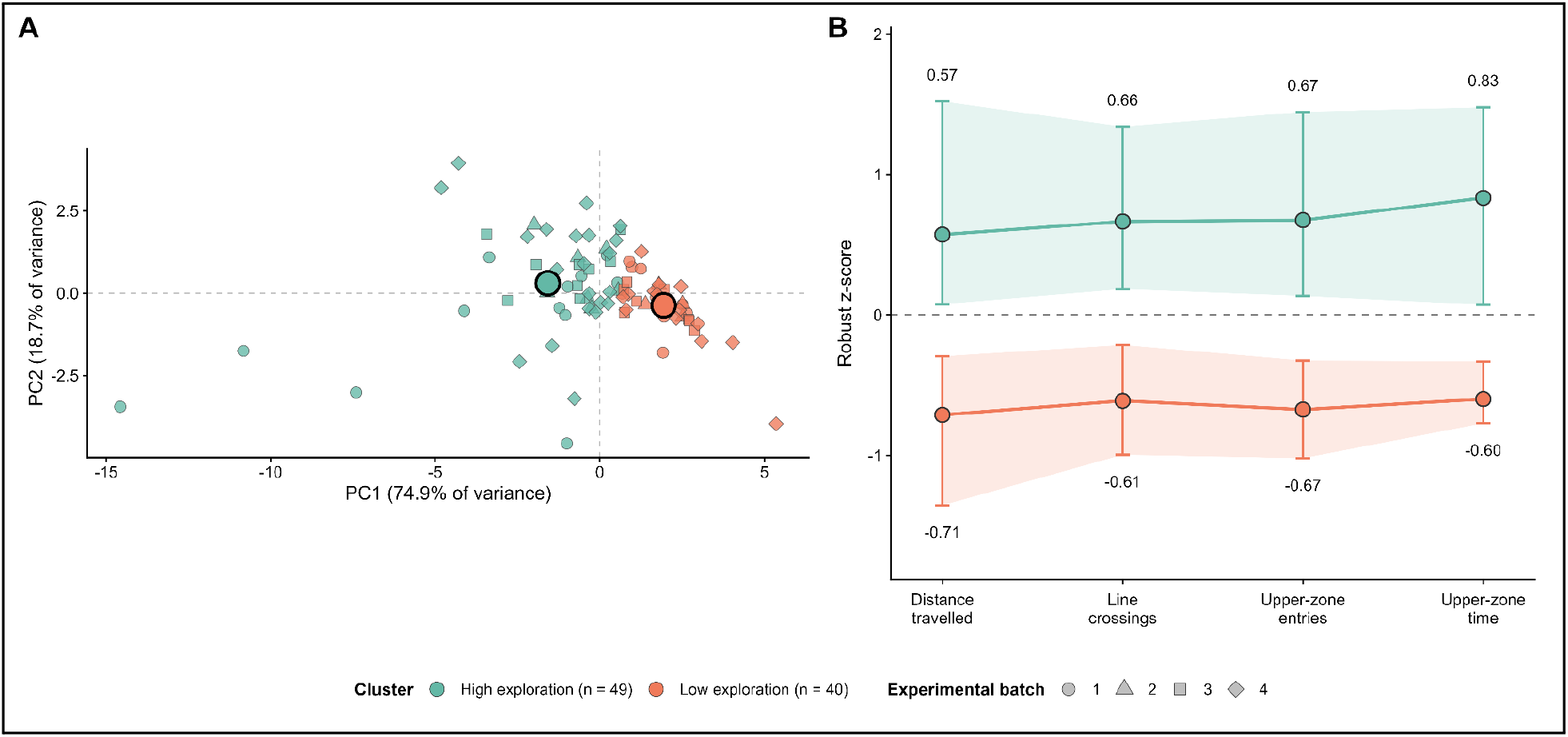
Novelty-evoked exploratory space use in the unstressed reference population forms a continuum with an operational two-profile partition. (A) Principal component analysis provided a two-dimensional representation of the control-referenced behavioural space for visualisation and was not used to derive the graph partition. PC1 and PC2 accounted for 74.9% and 18.7% of the total variance, respectively. Colours denote the two-cluster partition identified using a weighted *k*-nearest-neighbour network and Louvain community detection at the representative configuration, *k* = 45; identical individual-level memberships were obtained across *k* = 44–46. The resulting clusters were subsequently characterised as higher-exploration (*n* = 49) and lower-exploration (*n* = 40) reference profiles within the continuum of novelty-evoked exploratory space use. Point shapes identify the four independent experimental cohorts, and large black-outlined circles indicate the profile centroids calculated from the mean principal-component scores. (B) Higher- and lower-exploration reference profiles across the four cohort-specific, control-referenced robust coordinates. Points and numerical labels represent the median control-referenced robust score for each profile, vertical bars indicate the interquartile range, and shaded ribbons connect the corresponding first and third quartiles across behavioural dimensions. The higher-reference profile showed consistently higher values for distance travelled, line crossings, upper-zone entries and upper-zone time. These differences are presented descriptively; no inferential tests or effect sizes were calculated because the same four variables were used to construct the behavioural graph and derive the operational two-profile partition.

The reference population comprised 89 zebrafish from the LAPCOM historical database, drawn from four independent experimental cohorts conducted at different times, under conditions previously described in Bertelli et al., 2021; Marcon et al., 2019; Neves et al., 2026 and Sachett et al., 2022, but without exposure to stressors. For each zebrafish, we analysed four behavioural variables: distance travelled, line crossings, entries into the upper zone, and time spent in the upper zone (Fig. S1A). Although all four variables were positively correlated, none was fully redundant, indicating that they captured partially distinct components of locomotor activity and vertical exploration and were therefore retained for the subsequent multivariate analyses (Fig. S1B). PCA of the four control-referenced robust coordinates showed that PC1 and PC2 explained 74.9% and 18.7% of the total variance, respectively, accounting jointly for 93.6%.

The broad inter-individual dispersion and heterogeneous distributional structure observed across the behavioural variables raised the possibility that the unstressed reference population might contain recurrent multivariate configurations of novelty-evoked exploratory space use rather than constitute a homogeneous behavioural population (Fig. 1B and Fig. S1A). To examine this possibility, we constructed weighted, undirected *k*-nearest-neighbour networks based on Euclidean distances calculated in the four-dimensional behavioural space and applied the Louvain community-detection algorithm to identify clusters of individuals with similar multivariate behavioural profiles. For detailed descriptions and applications of this graph-based clustering approach, see Blondel et al., 2008 and Rajput et al., 2022.

We systematically evaluated neighbourhood sizes from *k* = 5 to *k* = 80 and characterised each resulting partition using the Calinski–Harabasz index, mean silhouette width and Davies–Bouldin index. Rather than selecting the isolated optimum of any single metric, we searched for regions in which the inferred cluster structure remained stable across consecutive values of *k* (Rajput et al., 2022). Both two- and three-cluster partitions emerged across the evaluated range. However, the two highest Calinski–Harabasz values, the two highest mean silhouette widths and the two lowest Davies–Bouldin values all corresponded to two-cluster solutions (Fig. S1C). An exact-equivalence plateau was identified from *k* = 44 to *k* = 46, with identical two-cluster memberships across all three neighbourhood sizes and an ARI of 1.000 between adjacent partitions. The central value, *k* = 45, was retained as the algorithmic representative for null-model and resampling analyses. At this value, the Calinski–Harabasz index was 37.908, the mean silhouette width was 0.296, and the Davies–Bouldin index was 1.039. For subsequent projection analyses, the identical partition obtained at *k* = 46 was retained as the authorised fixed configuration; because *k* = 44–46 produced identical individual memberships, this choice did not alter the underlying two-profile partition.

To determine whether the resulting clusters represented genuinely discrete behavioural communities, the observed weighted modularity was compared with that obtained from 1,000 Gaussian-copula null simulations exactly preserving the empirical marginal distribution of each behavioural variable while approximately retaining their rank-dependence structure. The observed modularity (*Q* = 0.217) was close to the null expectation (mean *Q_null_* = 0.207, *SD* = 0.077; 2.5^th^–97.5^th^ percentiles = 0.092–0.440) and was not significantly greater than expected after accounting for the selection of *k* (*z* = 0.139, empirical *p* = 0.393); the fixed-*k* sensitivity analysis produced the same conclusion (*z* = −0.210, *p* = 0.592). Thus, the two-cluster partition showed no detectable excess modular segregation beyond that expected from continuous, correlated behavioural variation.

Bootstrap resampling further revealed uncertainty in both membership and partition granularity. Across 1,000 iterations, agreement with the full-sample partition, calculated over unique original individuals, was appreciable but variable (mean ARI = 0.740, *SD* = 0.142; median ARI = 0.738, *IQR* = 0.631–0.855; 2.5^th^–97.5^th^ percentiles = 0.475–1.000). Two clusters were recovered in 40.8% of iterations, whereas three clusters were recovered most frequently (52.9%) and four clusters occurred in 6.3%. A *post hoc* sensitivity analysis based on 1,000 stratified 80% subsamples without replacement (*n* = 71) additionally demonstrated dependence on the relationship between neighbourhood size and sample size. When *k* was held at 45, mean agreement was ARI = 0.692 , and two-, three-, and four-cluster solutions were recovered in 34.0%, 61.5%, and 4.5% of subsamples, respectively. When *k* was reduced proportionally to 36, mean agreement increased to ARI = 0.864, and the corresponding recovery rates were 75.0%, 24.8%, and 0.2%. The marked change in cluster recovery following this adjustment indicates that partition granularity was sensitive to the neighbourhood size used in graph construction.

Taken together, the internal validation indices supported retaining the two-cluster solution as an operational representation of the complete sample, but neither the modularity test nor the resampling analyses supported its interpretation as two invariant, naturally discrete populations. Instead, the results are more consistent with a broad, resolution-dependent operational bipartition within a continuous behavioural space, whose precise subdivision depends partly on sample composition and graph resolution.

The resulting organisation was consistent with higher- and lower-exploration regions within a continuum of novelty-evoked exploratory space use (Fig. 1A) and was reminiscent of the bold–shy axis described in the neuroethological literature (Beigloo et al., 2024; Espigares et al., 2025; Fernando et al., 2024; Fontana et al., 2025; Hubená et al., 2026). However, because the present analysis was restricted to four variables measured within a single behavioural context, these regions were not interpreted as definitive bold and shy personality types. Instead, they were characterised as higher- and lower-exploration reference profiles within the continuum of novelty-evoked exploratory space use. The higher-exploration reference profile comprised 49 zebrafish (55.1%), whereas the lower-exploration reference profile comprised 40 zebrafish (44.9%; Fig. 1A). Zebrafish assigned to the higher-exploration profile showed greater distance travelled (median robust *z*-score: 0.572 versus −0.712), more line crossings (0.665 versus −0.612), more frequent entries into the upper zone (0.674 versus −0.674), and more time spent in the upper zone (0.833 versus −0.599; Fig. 1B). Because these variables were also used to construct the behavioural graph, the corresponding differences were interpreted exclusively as a descriptive characterisation of the profiles and were not subjected to inferential testing or regarded as independent validation of the partition.

Collectively, in the absence of UCS, inter-individual variation in novelty-evoked exploratory space use within the reference population was organised predominantly along a locomotor–exploratory continuum, with two broad profiles occupying opposite regions of this behavioural space: one characterised by greater locomotor activity and greater use of the upper zone, and the other by lower expression of these same behavioural dimensions. Thus, although the two-profile solution provided the most internally supported representation of the reference population, these profiles were interpreted as operational regions within a continuous behavioural space rather than as discrete behavioural populations.

### 3.2 Unpredictable chronic stress redistributes novelty-evoked exploratory space use within the control-derived behavioural reference architecture

To determine whether UCS redistributed novelty-evoked exploratory space use relative to the control-derived behavioural reference architecture described above, we projected 89 UCS-exposed zebrafish from the LAPCOM historical database into the four-dimensional control-referenced behavioural space defined by distance travelled, line crossings, upper-zone entries and upper-zone time (Fig. 2). The UCS-exposed population was significantly displaced relative to the control population (Fig. 2A; PERMANOVA: *F*(1,173) = 8.10; *R^2^*= 0.0423; *p* = 0.0016) towards lower locomotor activity and reduced vertical exploration (*D*₄D = 1.264).

**Figure 2.**
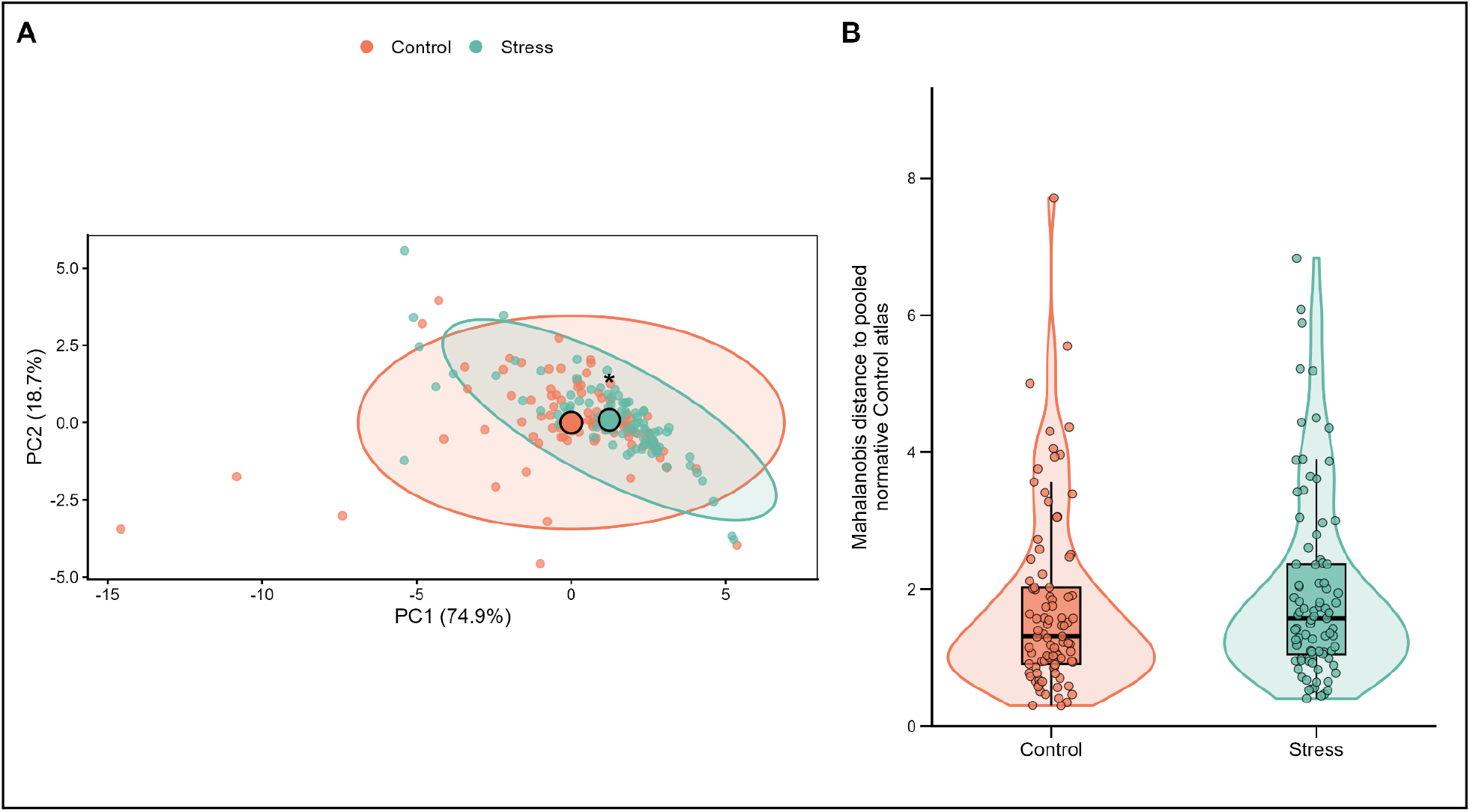
Unpredictable chronic stress redistributes novelty-evoked exploratory space use within the control-derived behavioural reference architecture. (A) Projection of 89 unstressed reference and 89 UCS-exposed zebrafish into the fixed PC1–PC2 space derived exclusively from the unstressed reference population. The PCA was fitted to the four cohort-specific, control-referenced robust behavioural coordinates obtained for distance travelled, line crossings, upper-zone entries and upper-zone time. UCS-exposed zebrafish were projected into this space using the fixed control PCA centre and rotation matrix. PC1 and PC2 explained 74.9% and 18.7% of the variance within the unstressed reference population, respectively. Each point represents one zebrafish; large black-outlined circles indicate the group centroids, and shaded contours represent descriptive 95% normal-theory data ellipses. The two independent populations remained extensively overlapping despite their centroidal displacement. The asterisk indicates a treatment-associated difference in multivariate location (PERMANOVA: *F*(1, 173) = 8.10, *R*² = 0.0423, *p* = 0.0016). The four-dimensional centroid displacement (*D*₄D), calculated as the Euclidean distance between the unstressed reference and UCS centroids in the complete four-dimensional robust behavioural space, was 1.264. (B) Distribution of individual square-root Mahalanobis distances from the control-derived behavioural reference distribution in 89 unstressed reference and 89 UCS-exposed zebrafish. For unstressed reference zebrafish, distances were calculated using leave-one-out estimates in which each individual was excluded from estimation of the reference centre and covariance matrix. Distances for UCS-exposed zebrafish were calculated relative to the complete control-derived reference centre and covariance matrix. Each point represents one zebrafish; violin contours show the estimated distributions, whereas the embedded boxplots indicate the median and interquartile range, with whiskers extending to the most extreme values within 1.5 times the interquartile range. Mahalanobis distances did not differ significantly between groups (median: unstressed reference = 1.31, UCS-exposed = 1.57; two-sided reference-reconstructing permutation test, *p* = 0.171; rank-biserial correlation, *r*ᵣᵦ = 0.119). Coral and turquoise represent unstressed reference and UCS-exposed zebrafish, respectively.

Upper-zone time contributed 58.6% of the squared centroidal displacement, followed by distance travelled (18.3%), upper-zone entries (15.2%), and line crossings (7.9%). Multivariate dispersion did not differ significantly between groups (PERMDISP: *F*(1,176) = 0.624; *p* = 0.420), although the UCS population distribution was descriptively more compact.

To determine how this displacement related to the control-derived reference profiles, each UCS-exposed zebrafish was assigned to the nearest centroid of the two control-derived profiles; no new *k*-nearest-neighbour graph or Louvain partition was generated for the UCS-exposed population. The proportion assigned to the higher-exploration profile was lower in the UCS-exposed group than in the control group (24.7% vs 55.1%; Fisher’s exact test, *p* = 5.98 × 10^−5^; Cramér’s *V* = 0.310). Nevertheless, the groups did not differ in covariance-adjusted Mahalanobis distance from the control-derived reference (primary reference-reconstructing permutation test: median difference UCS − control = 0.260, rank-biserial correlation (*r*ᵣᵦ) = 0.119, *p* = 0.171; secondary Mann–Whitney *U* test: *U*_control_ = 3490, *p* = 0.171; Fig. 2B) or in Euclidean distance to the nearest reference-profile centroid (Mann–Whitney *U* test : *U*_control_ = 4221, *r*ᵣᵦ = −0.0658, *p* = 0.449). The global correlation structure likewise showed no detectable reorganisation (unique-edge *L_2_*= 0.231, RMS Δ*r_s_* = 0.0945, maximum ∣Δ*r_s_*∣ = 0.127, permutation *p* = 0.325). The descriptive 95% normal-theory data ellipses showed extensive overlap (Fig. 2A). Thus, UCS primarily redistributed individuals towards the lower-exploration region of the control-derived behavioural reference architecture, with no detectable evidence that UCS produced configurations of novelty-evoked exploratory space use extending beyond the behavioural range represented by the unstressed reference population.

### 3.3 *De novo* post-UCS clusters of novelty-evoked exploratory space use show asymmetric correspondence with control-derived reference profiles

Building on the control-derived behavioural reference architecture established in the unstressed population, we next asked whether inter-individual heterogeneity in post-UCS novelty-evoked exploratory space use exhibited recurrent multivariate configurations, and whether these configurations, once interpreted relative to the control-derived reference profiles, could contribute to the operational identification of resilient- and susceptible-like archetypes. To characterise the internal organisation of the UCS-exposed population without imposing the control-derived profile assignments, the 89 UCS-exposed zebrafish were first analysed *de novo* in the four-dimensional control-referenced robust behavioural space (Fig. 3 and Fig. S2A–B). PCA revealed that PC1 accounted for 80.2% of the variance and PC2 for a further 13.3% (93.5% cumulative), with positive PC1 loadings for all four behavioural variables, indicating that the dominant dimension of post-UCS variation represented a general exploratory gradient (Fig. 3A).

**Figure 3.**
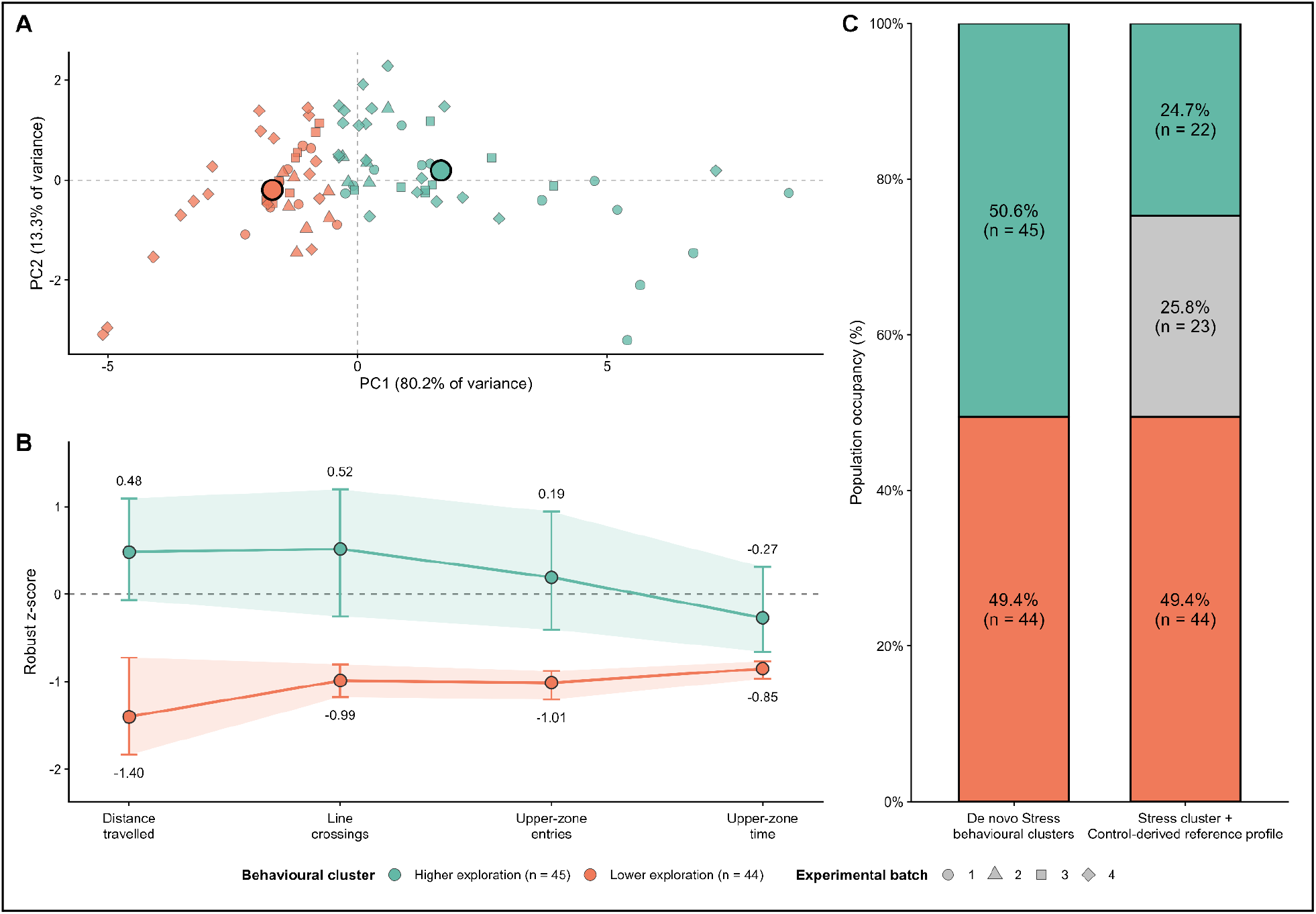
*De novo* post-UCS clusters of novelty-evoked exploratory space use show asymmetric correspondence with control-derived reference profiles. (A) *De novo* organisation of the UCS-exposed population (*n* = 89) within the complete four-dimensional control-referenced robust behavioural space. PCA provided a two-dimensional representation for visualisation and was not used to derive the cluster partition. PC1 and PC2 explained 80.2% and 13.3% of the variance within the UCS-exposed population, respectively. Clusters were identified directly in the complete four-dimensional space using a weighted, undirected union-*k*-nearest-neighbour graph and Louvain community detection at *k* = 48. Individual zebrafish are coloured according to the resulting higher-exploration (*n* = 45) and lower-exploration (*n* = 44) post-UCS clusters; point shapes denote the four experimental cohorts, and large black-outlined circles indicate the cluster centroids calculated from the mean principal-component scores. (B) Descriptive characterisation of the two *de novo* post-UCS clusters across the four control-referenced robust behavioural coordinates. Points and connecting lines represent cluster medians, vertical intervals and shaded regions represent the interquartile range, and numerical labels indicate the median control-referenced robust score for each variable. Turquoise and coral denote the higher- and lower-exploration clusters, respectively. No inferential comparisons are shown because these variables contributed directly to the derivation of the clusters. (C) Integration of *de novo* post-UCS cluster membership with assignment to the control-derived reference profiles. The initial *de novo* partition comprised 45 higher-exploration zebrafish (50.6%) and 44 lower-exploration zebrafish (49.4%). All 44 zebrafish belonging to the lower-exploration cluster were also assigned to the lower-exploration control-derived reference profile and were therefore operationally classified as susceptible-like archetypes. In contrast, the higher-exploration cluster showed incomplete correspondence with the control-derived reference architecture: 22 of 45 zebrafish were assigned to the higher-exploration reference profile and were operationally classified as resilient-like archetypes (24.7% of the UCS-exposed population), whereas the remaining 23 were assigned to the lower-exploration reference profile and remained unresolved (25.8%). Thus, membership in the higher-exploration post-UCS cluster alone was insufficient to define a resilient-like archetype. Turquoise, coral and grey denote resilient-like, susceptible-like and unresolved classifications, respectively. These designations represent operational classifications based on novelty-evoked exploratory space use and were not interpreted as independently validated biological phenotypes.

Internal structure within this space was then examined using a weighted, undirected union-*k*NN graph constructed from Euclidean distances and Louvain community detection (Fig. S2).

Candidate solutions were evaluated across *k* = 5–80. A stable two-cluster plateau extended from *k* = 41 to 56, with *k* = 48 selected as the representative partition (Fig. S2C). At *k* = 48, clustering quality was characterised by a mean silhouette width of 0.334, a Calinski–Harabasz index of 55.034 and a Davies–Bouldin index of 0.971. The partition comprised a behavioural cluster characterised by lower exploration, comprising 44 zebrafish (49.4%) and a behavioural cluster characterised by higher exploration, comprising 45 zebrafish (50.6%; Fig. 3A). Algorithmic stability was nearly complete: 499 of 500 Louvain repetitions reproduced the retained partition exactly, all 500 recovered two clusters, and the mean pairwise ARI was 0.999. Sampling stability was lower, with a mean bootstrap ARI of 0.806, a median of 0.826, and recovery of the two-cluster solution in 547 of 1,000 bootstrap iterations (54.7%); three clusters emerged in 44.4% and four in 0.9% of iterations.

The two behavioural clusters represented markedly different exploratory configurations (Fig. 3B). The lower-exploration cluster showed mean robust *z*-score of −1.55 for distance travelled, −1.01 for line crossings, −1.06 for upper-zone entries, and −0.828 for upper-zone time, whereas the corresponding values in the higher-exploration cluster were 0.608, 0.786, 0.642, and 0.0636.

The Euclidean distance between cluster centroids was 3.402 (Fig. 3A). Descriptive between-cluster contrasts were large for all four variables, with *r*ᵣᵦ of −0.906 for distance travelled, −0.851 for line crossings, −0.835 for upper-zone entries, and −0.694 for upper-zone time. Because these variables were used to derive the clusters, the corresponding effect sizes were interpreted exclusively as descriptive characterisations of the retained partition and not as independent validation.

We next asked whether these *de novo* post-UCS behavioural clusters corresponded to the control-derived reference profiles (Fig. 3C). The correspondence was asymmetric and incomplete (ARI = 0.227; individual-level directional agreement = 74.2%, 66/89 zebrafish). All 44 zebrafish in the lower-exploration behavioural cluster were assigned to the lower-exploration reference profile (44/44, 100%), thereby satisfying both behavioural criteria for their operational classification as susceptible-like archetype. In contrast, the higher-exploration behavioural cluster was heterogeneous: 22 of 45 zebrafish (48.9%) were assigned to the higher-exploration reference profile, whereas 23 of 45 (51.1%) were still assigned to the lower-exploration reference profile. Thus, membership in the higher-exploration behavioural cluster alone was insufficient to define resilience-like archetype.

We therefore operationally classified as resilient-like archetype the 22 zebrafish that satisfied both criteria: membership in the behavioural cluster characterised by higher exploration that emerged *de novo* after UCS and assignment to the higher-exploration control-derived reference profile. These animals represented 24.7% of the UCS-exposed population and comprised all UCS-exposed zebrafish assigned to the higher-exploration reference profile in the preceding population-level analysis. Collectively, these findings indicate that resilient-like responses to UCS did not emerge as a simple counterpart of susceptibility or as a discrete behavioural community identified through graph-based community detection. Rather, resilient-like archetypes were identified within a subset of the higher-exploration post-UCS configuration and were most stringently operationalised by requiring concordance between *de novo* organisation of novelty-evoked exploratory space use within the UCS-exposed population and assignment to the higher-exploration profile of the control-derived behavioural reference architecture.

### 3.4 Operational classification of post-UCS resilient- and susceptible-like archetypes is method-dependent but shows partial cross-method consensus

The internal organisation of novelty-evoked exploratory space use within the UCS-exposed population revealed a methodological challenge in the operational identification of resilient- and susceptible-like archetypes. Membership in a *de novo* post-UCS behavioural configuration was not, by itself, sufficient to establish a resilient-like classification, because its interpretation depended on the relationship between post-UCS organisation and the control-derived behavioural reference architecture (Fig. 3). This raised a broader methodological question: whether distinct analytical frameworks designed to identify multivariate behavioural configurations or latent population structure would produce comparable partitions when applied to the same post-UCS novelty-evoked exploratory space-use data, and whether those partitions would converge in their operational identification of resilient- and susceptible-like archetypes. To address this question, five analytical classification frameworks representing distinct strategies for partitioning behavioural variation were applied to the same 89 UCS-exposed zebrafish: a Swaminathan-like procedure specifically developed to distinguish between resilient and susceptible stress responses in zebrafish larvae by applying an optimal threshold to a continuous BI (Swaminathan et al., 2023); a PCA-derived index quantile classification that defined susceptible, intermediate, and resilient stress response groups from the lower, middle, and upper portions of that index (Espigares et al., 2025; Leong et al., 2025; Yata and Aoshima, 2020); Gaussian mixture modelling, which identified latent probabilistic subgroups in multivariate behavioural space (Fraley and Raftery, 2007; Videlier et al., 2015); Two-Step clustering, which partitioned individuals according to multivariate similarity using an automated model-selection procedure (Benassi et al., 2020; IBM Corp., 2025; Mueller et al., 2021); and Louvain community detection, which identified densely connected groups within the behavioural similarity network (Beigloo et al., 2024; Blondel et al., 2008; Rajput et al., 2022).

To provide a common directional framework for interpreting the resulting classes, each zebrafish was additionally characterised by a BI derived from its scores across the four control-derived PCA axes and oriented such that higher values indicated greater alignment with the control-like behavioural direction, whereas lower values denoted greater displacement towards the UCS-associated configuration. The BI served as the direct classification variable for the Swaminathan-like and PCA-derived index quantile procedures, whereas for GMM, Two-Step clustering, and Louvain community detection it was not used to generate the partitions, but rather to orient and compare the resulting classes along the same directional behavioural scale. The mean BI therefore represents the average position of the zebrafish assigned to a given class along this common behavioural continuum, allowing classes generated by analytically distinct methods to be compared directly. Because the BI was derived from the same four behavioural variables, its use for class orientation was descriptive and did not provide independent validation of the resulting classifications.

The five approaches generated markedly different population partitions, both in the number of categories recovered and in the proportion of zebrafish assigned to each operational class (Fig. 4 and Table 1). Among the binary procedures, the Swaminathan-like threshold classified 24 of 89 zebrafish as resilient (27.0%) and 65 as susceptible (73.0%), while Two-Step clustering yielded a closely comparable distribution of 21 resilient (23.6%) and 68 susceptible individuals (76.4%). For the method-comparison analysis, the two Louvain communities were ordered according to their mean BI and operationally labelled resilient-like archetype (45 zebrafish, 50.6%) and susceptible-like archetype (44 zebrafish, 49.4%). These method-specific labels were distinct from the stricter reference-integrated classification described above. The two approaches retaining an explicit intermediate category generated qualitatively different distributions. PCA-derived index quantile classification classified 23 zebrafish as resilient (25.8%), 43 as intermediate (48.3%), and 23 as susceptible (25.8%), whereas GMM assigned only 9 zebrafish to the resilient class (10.1%), 52 to the intermediate class (58.4%), and 28 to the susceptible class (31.5%). Consequently, across the five operational definitions, the number of zebrafish classified as resilient ranged from 9 to 45, while susceptible classifications ranged from 23 to 68 individuals (Table 1). Thus, even before considering whether the same individuals were assigned to corresponding classes, the proportion of zebrafish operationally classified as resilient or susceptible varied substantially according to the analytical framework employed.

**Figure 4.**
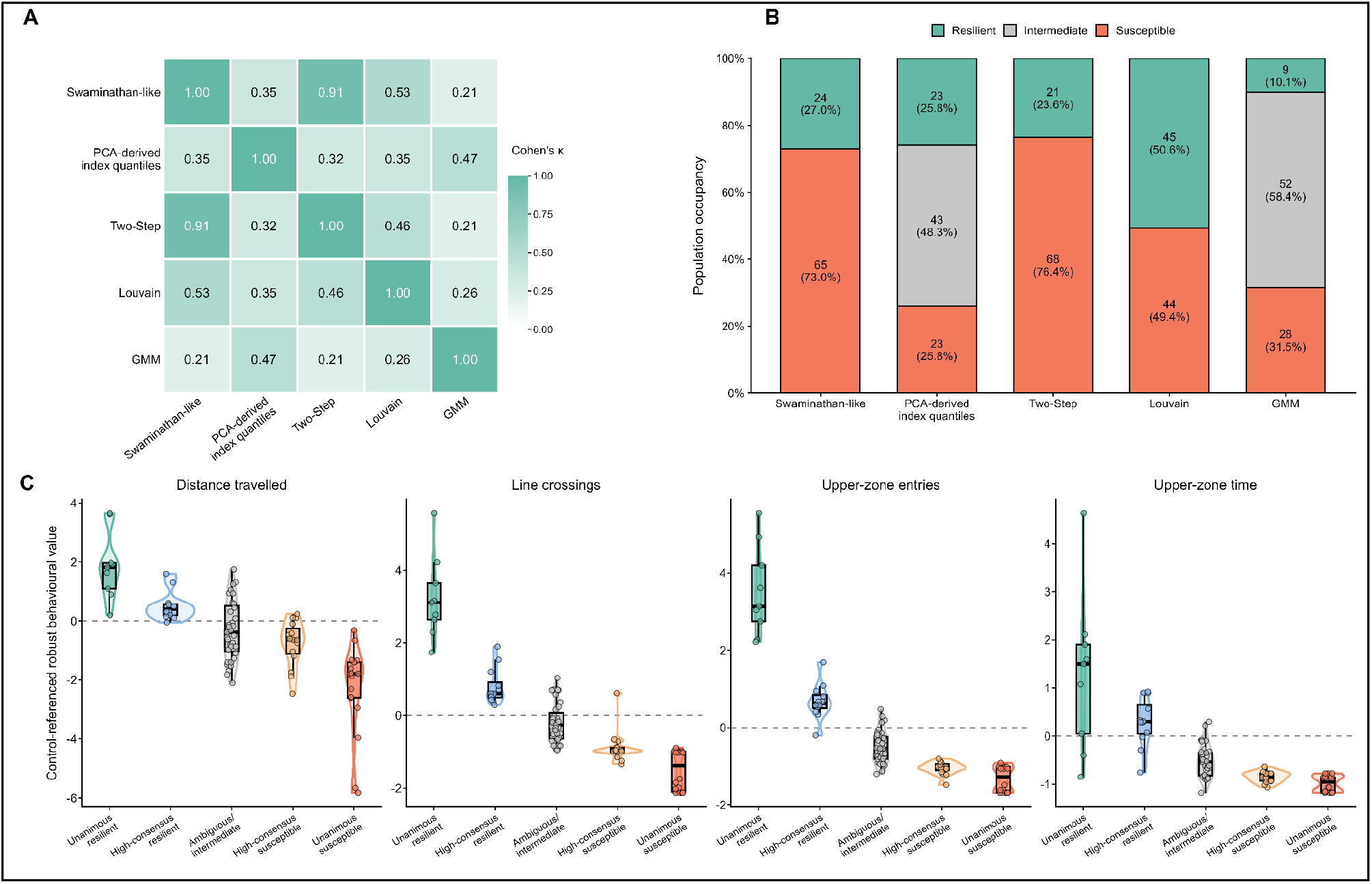
Alternative analytical frameworks yield method-dependent classifications of post-UCS behavioural variation with partial individual-level convergence. (A) Pairwise agreement among the five analytical classification frameworks, quantified using Cohen’s *κ* to account for agreement expected by chance. Cell values and colour intensity represent agreement in individual-level assignments across the same 89 UCS-exposed zebrafish. Because the frameworks differ in both their category structure and operational definitions, these comparisons quantify concordance between classifications rather than equivalence of the underlying constructs. (B) Relative population occupancy under each analytical framework, showing the proportion of UCS-exposed zebrafish assigned to the corresponding method-specific categories. Swaminathan-like thresholding classified individuals as resilient or susceptible stress responses; Two-Step clustering yielded a binary resilient/susceptible classification; Louvain community detection produced two *de novo* behavioural communities that, for cross-method comparison only, were ordered according to their mean Behavioural Index and operationally labelled resilient-like and susceptible-like. PCA-derived index quantile classification and Gaussian mixture modelling yielded three-category solutions including an intermediate class. The Louvain labels shown here are method-specific and are distinct from the stricter reference-integrated resilient- and susceptible-like archetypes defined in Figure 3. (C) Distributions of the four control-referenced robust behavioural scores across the five cross-method consensus categories. For cross-method consensus, corresponding extreme categories were directionally aligned across analytical frameworks without implying equivalence of their underlying definitions. Unanimous categories comprised zebrafish assigned to the corresponding extreme by all five frameworks, whereas high-consensus categories comprised those receiving the corresponding assignment from four of the five frameworks. Zebrafish not meeting either high-consensus criterion were retained in the lower-consensus group. Violin contours show the distributions of individual observations; embedded boxplots indicate the median and interquartile range, and individual zebrafish are shown as jittered points. Dashed horizontal lines denote the control-referenced value of zero. Descriptive effect sizes across consensus categories were large for all four behavioural variables (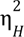 = 0.556–0.810). Because the same behavioural information contributed to the construction or orientation of the underlying classifications, these contrasts were interpreted exclusively as descriptive characterisations of cross-method consensus and not as independent biological validation.

**Table 1.** Alternative analytical frameworks yield method-dependent operational classifications of post-UCS behavioural variation. The table summarises the number and percentage of UCS-exposed zebrafish (*n* = 89) assigned to the method-specific categories generated or operationally interpreted under each analytical framework. Swaminathan-like thresholding classified individuals as resilient or susceptible stress responses, following the conceptual framework of the original procedure. PCA-derived index quantile classification defined resilient, intermediate and susceptible stress-response groups, whereas Gaussian mixture modelling identified three latent probabilistic classes that were subsequently oriented along the control-referenced behavioural continuum. Two-Step clustering generated a binary multivariate partition whose classes were directionally interpreted as resilient and susceptible for method comparison. Louvain community detection identified two *de novo* behavioural communities; for the cross-method comparison only, these were ordered according to their mean Behavioural Index and operationally labelled resilient-like and susceptible-like. These Louvain labels were distinct from the stricter reference-integrated resilient- and susceptible-like archetypes defined in Figure 3. Swaminathan-like thresholding, Two-Step clustering and Louvain community detection therefore yielded binary classifications, whereas PCA-derived index quantile classification and Gaussian mixture modelling retained an intermediate category. The corresponding individual-level classifications are shown in Figure 4B. Values are presented as *n* (%); dashes indicate that the corresponding analytical framework did not define an intermediate category.

| **Statistical method** | **Resilient-like archetype, *n* (%)** | **Intermediate archetype, *n* (%)** | **Susceptible-like archetype, *n* (%)** | **Classification structure** |
|---|---|---|---|---|
| Swaminathan-like threshold | 24 (27.0%) | — | 65 (73.0%) | Binary |
| PCA-derived index quantile classification | 23 (25.8%) | 43 (48.3%) | 23 (25.8%) | Three-class |
| Gaussian mixture modelling | 9 (10.1%) | 52 (58.4%) | 28 (31.5%) | Three-class |
| Two-Step clustering | 21 (23.6%) | — | 68 (76.4%) | Binary |
| Louvain community detection | 45 (50.6%) | — | 44 (49.4%) | Binary |

This variation in class prevalence was accompanied by substantial differences in individual-level assignment (Fig. 4A–B and Fig. S3). The strongest convergence occurred between the Swaminathan-like threshold and Two-Step clustering, which assigned the same operational label to 86 of 89 zebrafish (96.6% exact agreement). Their classifications showed very high chance-corrected agreement (Cohen’s *κ* = 0.911) and strong similarity in partition structure (ARI = 0.861), with a strong association between classifications (Cramér’s *V* = 0.915; Fisher’s exact p = 1.55 × 10⁻¹⁷). Concordance decreased when either binary approach was compared with Louvain community detection. Swaminathan-like threshold and Louvain community detection agreed for 68 zebrafish (76.4%), with *κ* = 0.531 and ARI = 0.272 (Cramér’s *V* = 0.601; Fisher’s exact *p* = 1.72 × 10⁻⁹), whereas Two-Step and Louvain community detection agreed for 65 zebrafish (73.0%), with *κ* = 0.464 and ARI = 0.206 (Cramér’s *V* = 0.550; Fisher’s exact *p* = 4.44 × 10⁻⁸). Among the approaches retaining an intermediate category, PCA-derived index quantile classification and GMM classified 60 of 89 fish identically (67.4% exact agreement); although agreement corrected for chance was moderate (*κ* = 0.466), similarity between the resulting partitions remained comparatively limited (ARI = 0.250; Cramér’s *V* = 0.570; Monte Carlo Fisher *p* ≤ 1.0 × 10⁻⁵). GMM showed particularly low concordance with the two binary classifications, agreeing with both Swaminathan-like threshold and Two-Step clustering for only 37 zebrafish (41.6%; *κ* = 0.214 and 0.206; ARI = 0.138 and 0.153, respectively). Thus, the analytical framework influenced not only the estimated prevalence of resilient-like and susceptible-like, but also which individual zebrafish were assigned to each operational category. For the method-comparison analysis, the two Louvain communities were ordered according to their mean BI and operationally labelled resilient-like (45 zebrafish, 50.6%) and susceptible-like (44 zebrafish, 49.4%). These method-specific labels were distinct from the stricter reference-integrated classification described above and comparisons between binary and three-category frameworks were necessarily influenced by their different category structures.

Despite differences in individual assignment, all method-specific classes were directionally ordered along the control-referenced behavioural continuum, as expected from their construction and BI-based orientation (Fig. 5). Across methods, susceptible-like classes consistently occupied the lower end of the behavioural space, whereas resilient-like classes showed higher exploratory values for distance travelled, line crossings, upper-zone entries and upper-zone time. The magnitude of these between-class contrasts was descriptively large across the five analytical frameworks, with 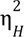 values ranging from 0.351 to 0.714. Descriptive separation was particularly pronounced for the PCA-derived index quantile classification, for which 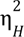 ranged from 0.579 for upper-zone time to 0.714 for upper-zone entries, and for GMM, which yielded 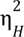 values ranging from 0.361 to 0.697 across the four variables. Among the binary approaches, rank-based effect sizes ranged from *r* = 0.645 to 0.742 for the Swaminathan-like threshold, from *r* = 0.595 to 0.722 for Two-Step clustering and from *r* = 0.598 to 0.780 for Louvain community detection. The two three-class procedures also placed the intermediate class between the resilient-like and susceptible-like extremes across the four behavioural dimensions, although upper-zone time showed comparatively weaker descriptive separation between the intermediate and resilient-like GMM classes. Because the same behavioural variables either directly informed classification or were incorporated into the BI used to classify or orient the resulting classes, these between-class contrasts were interpreted exclusively as descriptive characterisations of the partitions and not as independent validation of the classifications.

**Figure 5.**
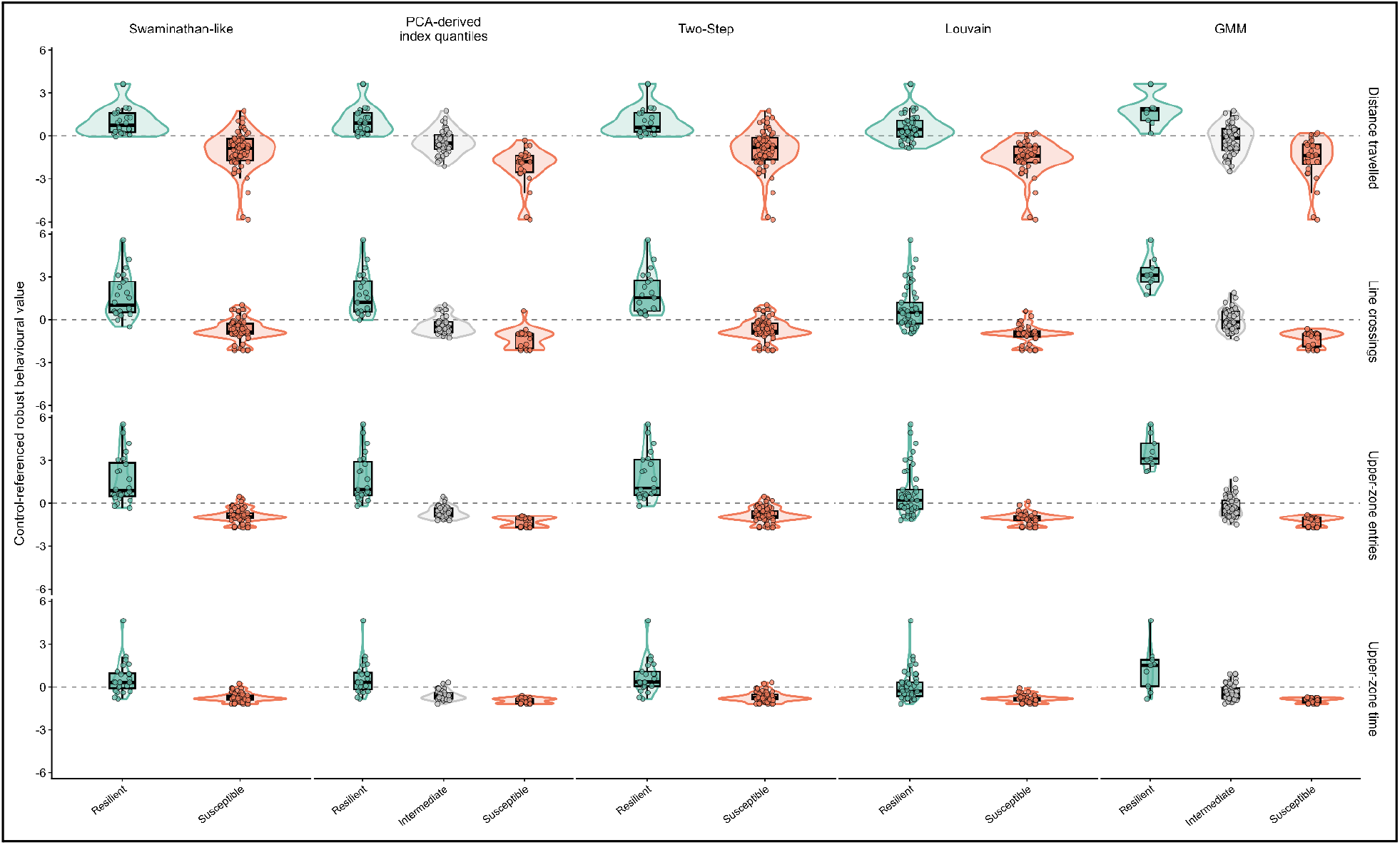
Behavioural distributions characterising the method-specific operational categories generated by alternative analytical frameworks. Distributions of the four control-referenced robust behavioural scores—distance travelled, line crossings, upper-zone entries and upper-zone time—across the method-specific categories generated by the five analytical frameworks. Columns correspond to the analytical frameworks and rows to the behavioural variables. For visual comparison across methods, categories occupying the higher, intermediate and lower portions of the control-referenced behavioural continuum are shown in turquoise, grey and coral, respectively; this common colour coding does not imply equivalence of the underlying operational definitions. Swaminathan-like thresholding classified zebrafish as resilient or susceptible stress responses, whereas Two-Step clustering yielded a binary multivariate classification. Louvain community detection generated two *de novo* behavioural communities that, for cross-method comparison only, were ordered according to their mean Behavioural Index and operationally labelled resilient-like and susceptible-like. PCA-derived index quantile classification and Gaussian mixture modelling yielded three-category solutions that included an intermediate category. Across methods and behavioural variables, descriptive between-category effect sizes ranged from 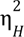 = 0.351 to 0.714. For the binary frameworks, descriptive contrasts between the two directionally opposed categories ranged from |*r*| = 0.595 to 0.780. Violin contours show the distributions of individual observations; embedded boxplots 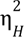 indicate the median and interquartile range, and individual zebrafish are shown as jittered points. Dashed horizontal lines denote the control-referenced value of zero. Because the same behavioural information contributed directly to the construction of some classifications or to the Behavioural Index used to orient others, the displayed contrasts and effect sizes were interpreted exclusively as descriptive characterisations of the resulting method-specific categories and not as independent biological validation.

Importantly, cross-method consensus revealed an ordered behavioural gradient (Fig. 3C and 5). Nine zebrafish were unanimously classified as resilient-like, while 12 additional zebrafish reached 4/5-method consensus for a resilient-like classification. At the opposite extreme, 18 zebrafish were unanimously classified as susceptible-like and an additional 15 reached 4/5-method consensus for a susceptible-like classification; the remaining 35 showed lower cross-method concordance. Mean BI values decreased progressively from 3.40 in unanimously resilient-like zebrafish to 0.254 in high-consensus resilient-like zebrafish, −1.46 in the lower-consensus group, −2.31 in high-consensus susceptible-like zebrafish, and −3.35 in unanimously susceptible-like zebrafish, accompanied by corresponding monotonic shifts across all four behavioural dimensions. Consensus-category differences were significant for distance travelled, line crossings, upper-zone entries, and upper-zone time (*p* = 2.61 × 10⁻¹⁰ to 8.47 × 10⁻¹⁵; 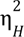 = 0.556–0.810). Collectively, these results show that alternative frameworks for partitioning behavioural variation do not recover an invariant population structure. Nevertheless, their partially overlapping classifications reveal a subset of zebrafish whose resilient-like or susceptible-like assignment remains consistent across markedly different analytical definitions of behavioural organisation.

## 4. Discussion

Taken together, the present findings indicate that novelty-evoked exploratory space use in adult zebrafish is organised predominantly as a continuous multivariate behavioural architecture rather than as a set of naturally discrete behavioural types. Within the unstressed reference population, graph-based community detection provided an operational two-profile representation of this continuum, but formal null-model inference and resampling did not support the interpretation of these profiles as invariant biological populations. UCS subsequently displaced the population within this pre-existing reference architecture towards lower locomotor activity and reduced vertical space use, while leaving substantial overlap with the unstressed reference population and without detectable evidence, within the four behavioural dimensions examined, of configurations extending beyond the reference range. *De novo* analysis of the UCS-exposed population nevertheless revealed marked inter-individual heterogeneity. Importantly, this heterogeneity was asymmetric with respect to the control-derived reference: all individuals in the lower-exploration post-UCS configuration also occupied the lower-exploration reference profile, whereas only 22 of the 45 individuals in the higher-exploration post-UCS configuration recovered the higher-exploration reference assignment. Higher exploratory expression after UCS was therefore insufficient, by itself, to define a resilient-like response. Finally, the estimated prevalence and individual identity of resilient- and susceptible-like classifications varied substantially across analytical frameworks, despite partial convergence among methods and the emergence of subsets with consistently extreme assignments. These results position behavioural heterogeneity itself as an important component of the post-UCS phenotype and indicate that its experimental stratification requires an explicit reference architecture and careful evaluation of the statistical structure used to generate classifications. Such stratification should be regarded as a phenotypic starting point for subsequent physiological and neurobiological investigation, rather than as evidence that resilience has already been biologically validated.

The continuous organisation of the unstressed population is consistent with an increasingly multidimensional view of zebrafish exploration. Rajput et al. showed, using three-dimensional trajectories from more than 400 adult zebrafish, that exploratory behaviour could not be reduced satisfactorily to a simple bold–shy dichotomy; unsupervised analysis recovered several behavioural configurations, and cluster membership was influenced by strain and sex. Moreover, only 54% of individuals retained the same cluster assignment across two consecutive exposures, although this exceeded permutation-based expectation, indicating that exploratory structure can coexist with considerable within-individual plasticity (Rajput et al., 2022). Their observations are particularly relevant to the present study because they distinguish the existence of structure in multivariate behavioural space from the stronger claim that such structure represents immutable behavioural types. Our results extend this distinction. Louvain community detection yielded a useful operational organisation of the unstressed behavioural space, yet the observed modularity was not greater than expected under a continuous correlated null architecture, and the number of communities showed sensitivity to resampling and graph resolution. Thus, the higher- and lower-exploration reference profiles should be understood as reproducible regions of a continuous behavioural landscape rather than evidence for two naturally discrete populations. This distinction is methodologically important because clustering algorithms necessarily return partitions when applied to appropriate data structures; the inferential question is whether the resulting partition contains biological information beyond that expected from continuous covariation alone.

The graph-based strategy used here was motivated by the capacity of network approaches to preserve local multivariate relationships without imposing an *a priori* behavioural threshold. A direct precedent in zebrafish exploratory behaviour was provided by Rajput et al. (Rajput et al., 2022), who constructed a *k*-nearest-neighbour network from four three-dimensional novel-tank behavioural measures and applied Louvain community detection to identify four behavioural clusters. The same group subsequently extended Louvain-based community detection to a distinct systems-level context, using correlations in *c-fos* activity among brain regions to identify community structure within a functional network associated with novel tank exposure (Rajput et al., 2025). Our implementation therefore did not introduce Louvain community detection into zebrafish behavioural phenotyping, but extended its inferential use by explicitly distinguishing the detection of an algorithmic partition from evidence that such a partition represents discrete population structure. Individual zebrafish constituted the network nodes, phenotypic similarity defined weighted neighbourhood relationships, and candidate partitions were evaluated across neighbourhood scales. Weighted modularity, formal null-model inference, bootstrap resampling and subsampling analyses were then used not simply to optimise a preferred partition, but to address progressively stronger questions about algorithmic reproducibility, sampling stability and evidence for behavioural discreteness. These properties are not interchangeable. In the present reference population, algorithmic reproducibility was strong, sampling stability was substantial but resolution-dependent, whereas evidence for genuinely discrete population structure was not supported. This distinction is particularly important when community detection is used upstream of biological classification, because an algorithmically reproducible partition can otherwise acquire biological meaning that the underlying data do not independently establish.

Against this reference architecture, UCS produced a statistically detectable but relatively modest multivariate displacement rather than a wholesale reorganisation of behavioural space. The treatment effect was significant in the partial PERMANOVA (*F*(1,173) = 8.10, *R²* = 0.0423, *p* = 0.0016), and the displacement of the UCS centroid reached *D_4_*D = 1.264. Consistent with this population-level shift, the proportion assigned to the higher-exploration reference profile decreased from 55.1% in the unstressed population to 24.7% after UCS. However, the two populations remained extensively overlapping; multivariate dispersion did not differ significantly, covariance-adjusted Mahalanobis distance from the control reference was not significantly increased, distance to the nearest control-derived centroid was unchanged, and the global Spearman correlation structure showed no detectable reorganisation. These convergent observations favour an interpretation in which UCS altered the relative occupancy of an existing behavioural architecture rather than creating, within the measured dimensions, an entirely distinct behavioural repertoire. This interpretation is intentionally narrower than the claim that UCS cannot generate qualitatively new behavioural states. The present behavioural space was defined by four cumulative variables, and behaviours absent from that space cannot be excluded simply because they were not observed as departures along those four axes.

That limitation is biologically consequential. Freezing frequency and duration, immobility, latency to first upper-zone entry, average vertical position, erratic swimming, turn angle, speed distributions, dwell times, transition rates, spatial coverage and path structure could reveal post-UCS configurations that are not reducible to quantitative displacement along the present locomotor–vertical gradient. This point is analogous to distinguishing severity along an existing dimension from the appearance of an additional behavioural dimension. The current finding that upper-zone time contributed 58.6% of the squared centroid displacement is particularly informative. It suggests that vertical organisation was a major component of the UCS-associated shift and supports prospective evaluation of average tank height as a continuous measure that may retain information lost when the water column is divided at an arbitrary boundary. Such measures may prove especially useful as candidate primary outcomes, although their prospective value should be established a priori rather than inferred from the same dataset in which they are selected. Three-dimensional tracking would further extend this approach. The present two-dimensional analysis captures vertical position effectively from the lateral plane but cannot reconstruct the complete volume explored, three-dimensional path geometry or spatial coverage that can be obtained from multi-camera approaches such as those used by (Rajput et al., 2022).

More fundamentally, cumulative six-minute endpoints collapse the temporal organisation of behaviour. Wong et al. demonstrated that transitions to the upper zone, time spent there, freezing and erratic movements change within a single novel-tank session, and that the greatest behavioural change in longer trials occurs during the early portion of exposure (Wong et al., 2010). Two individuals can therefore reach nearly identical six-minute totals through markedly different trajectories: rapid vertical expansion followed by stable occupancy, delayed ascent, repeated transitions between zones, prolonged initial immobility followed by exploration, or persistent oscillation. These trajectories need not represent equivalent behavioural states.

Future analyses should consequently preserve the serial structure of the NTT, whether through minute-wise bins or finer time series, and quantify latencies, slopes, dwell durations, transition probabilities and, where justified, behavioural change points or trajectory classes. Dunn et al. provide a neurobiological precedent for treating exploratory movement as sequentially organised rather than as an accumulation of independent motor events. In larval zebrafish, turn direction depended on recent turn history, and perturbation of the anterior rhombencephalic turning region altered both direction bias and this temporal dependence (Dunn et al., 2016). Although those findings cannot be transferred directly to adult NTT circuitry, they demonstrate a general principle: behavioural organisation resides not only in how much an animal moves, but also in the temporal relationships among successive actions.

The same reasoning applies spatially. Zebrafish establish homebases in novelty paradigms, repeatedly occupying and returning to particular locations rather than sampling the arena uniformly (Stewart et al., 2010). Thus, vertical occupancy, spatial transitions, homebase organisation and time-dependent familiarisation are related but non-equivalent components of the NTT. Wong et al. interpreted progressive increases in exploration and reductions in freezing as intra-session habituation to novelty, and pharmacological or alarm-cue manipulations altered these dynamics (Wong et al., 2010). In the present context, however, “habituation” should be used cautiously. Our cumulative historical data do not establish learning or memory updating at the individual level; a more conservative description is time-dependent behavioural reorganisation during continued exposure to novelty. This distinction matters because a six-minute NTT is not simply a brief measure of an enduring trait. It is an evolving interaction between an individual and an unfamiliar environment.

This also places limits on interpreting the NTT as a measure of motivation. Exposure to novelty is imposed: the fish is transferred to the unfamiliar tank rather than choosing whether to enter it. Once there, however, the animal must continuously select among alternative actions—remaining stationary, swimming, ascending, descending, changing direction or continuing to sample the environment. Novelty-evoked exploratory space use may therefore recruit neural systems involved in motivated action selection without constituting a direct measure of the motivation to seek novelty. This distinction is compatible with neurobiological evidence from zebrafish. Trotha et al. showed that pallial territories, particularly Dm, are recruited during behaviours involving stimulus valuation and motivated approach, including drug-seeking, while related territories are also engaged during aversive behaviour (Trotha et al., 2014). A stronger assay of exploratory motivation would require greater agency, for example voluntary entry from a familiar refuge into a novel compartment, repeated choice between familiar and novel environments, or willingness to incur a modest cost to access novelty. Such paradigms would complement rather than replace the NTT, allowing the motivation to engage with novelty to be separated from the organisation of behaviour after novelty has been imposed.

The ecological context in which these responses evolved should also temper simple interpretations of NTT variables. Field observations indicate that wild zebrafish inhabit shallow floodplain environments, including ponds and ditches with relatively clear water and marginal vegetation, and that they occupy the entire water column rather than exhibiting an obligatory preference for the bottom (Spence et al., 2006). Consequently, bottom restriction in a novel laboratory tank should not be assumed to reflect an intrinsic habitat preference. At the same time, natural risk contingencies are largely absent from conventional laboratory housing. Alarm substance exposure increases erratic swimming and other defensive responses in zebrafish (Speedie and Gerlai, 2008), whereas presentation of an animated predator reduces swimming velocity while increasing erratic movement, jumping and turn-angle measures (Gerlai et al., 2009). These observations caution against interpreting distance travelled, bottom dwelling, freezing or erratic swimming along a single anxiety axis. A reduction in mean locomotion can coexist with intense episodic defensive movement, and behaviour that appears maladaptive in a bare tank may have a different functional meaning in a natural setting containing substrate, vegetation and refuge opportunities. The proposal that bottom-oriented erratic movement might disturb sediment and contribute to concealment remains an ecologically plausible hypothesis rather than a demonstrated function in wild zebrafish; its importance here is conceptual, because it illustrates how behavioural meaning depends on environmental affordances.

Laboratory conditions therefore constitute part of the phenotype being measured. Tank shape, water depth, illumination, water source, visual enclosure, substrate, handling and previous social experience may all alter the information available to the fish and the actions that can be expressed (Golushko et al., 2025). This is relevant to the widespread use of both trapezoidal and rectangular novel tanks. In a trapezoidal apparatus, the horizontal cross-sectional area changes with height, geometrically coupling vertical position with the amount of available space (Wong et al., 2010); in a rectangular tank these relationships differ (Gallas-Lopes et al., 2023). Neither geometry is intrinsically superior, but they should not be assumed to instantiate identical behavioural problems. Wong et al., for example, used a 1.5-L trapezoidal tank and simultaneously quantified upper-zone occupancy, transitions, erratic movements and freezing (Wong et al., 2010). Environmental history is similarly relevant. Environmental enrichment can alter the response to UCS, including attenuation of UCS-associated cortisol and oxidative changes, indicating that enrichment is not simply a neutral acclimatisation procedure but a biological manipulation capable of modifying stress susceptibility (Marcon et al., 2018a). Introducing enrichment before UCS would therefore require a newly defined reference architecture rather than being treated as a procedural refinement without phenotypic consequences.

The neural organisation of the NTT further argues against reducing the assay to a single behavioural construct. Whole-brain mapping in adult zebrafish identified widespread *c-fos* recruitment after novel-tank exposure, including pallial, subpallial, thalamic, hypothalamic, cerebellar and neurotransmitter-associated territories (Rajput et al., 2025). Network analysis highlighted the ventral telencephalon, preoptic regions, Vmn and olfactory bulb as prominent components, with internal and external olfactory-bulb layers occupying high-centrality positions (Rajput et al., 2025). This *c-fos*-derived network should not be interpreted as an anatomical or causal connectome, but it supports the view that novel-tank behaviour emerges from distributed integration of sensory assessment, internal state and action selection rather than from a single “anxiety centre”. Notably, global *c-fos* expression peaked approximately 15 min after behaviour and then declined, emphasising that endpoint molecular mapping and within-test behavioural dynamics operate on different temporal scales (Rajput et al., 2025). Rajput et al. detected a strong temporal effect on *c-fos*, only a small non-significant trend for sex and no sex-by-time interaction in that experiment; this should not be generalised to the absence of sex effects on exploratory behaviour itself, particularly given established sex- and strain-related differences in behavioural clustering (Rajput et al., 2025, 2022). Sex, source tank and rack therefore remain appropriate blocking factors in prospective designs, especially where transfer and regrouping may modify social hierarchies before testing.

The asymmetry observed in our *de novo* post-UCS analysis is particularly important for the interpretation of resilience. All 44 fish belonging to the lower-exploration post-UCS configuration also mapped to the lower-exploration control reference profile, providing a convergent operational basis for a susceptible-like designation. The reverse was not true for the higher-exploration configuration: only 22 of 45 fish mapped to the higher-exploration reference profile, whereas the remaining 23 still occupied the lower-exploration reference region. Thus, more exploration after UCS cannot be equated with behavioural resilience. This finding also illustrates why a contemporaneous unstressed cohort is indispensable. Without the control-derived reference architecture, the 45 higher-exploration UCS fish could have been treated as a homogeneous “resilient” group despite their markedly different relationship to the reference population. At the same time, the use of a population reference does not establish the pre-UCS state of each individual. We cannot infer that post-UCS position reflects a fish’s premorbid “personality”, because the same individuals were not behaviourally phenotyped before stress. A future longitudinal design could address this issue through non-invasive baseline measurements in the home tank or another paradigm that does not compromise subsequent novelty testing.

The comparison among analytical frameworks reinforces the same caution from a different direction. The number of individuals identified at the resilient extreme ranged from 9 to 45 across methods, while susceptible classifications ranged from 23 to 68. Even the strongest agreement—between the Swaminathan-like threshold and Two-Step classification—does not constitute independent validation, because both operate on information derived from the same behavioural domain. Likewise, the unanimous and high-consensus subsets are scientifically useful precisely because their classifications are comparatively insensitive to the operational rule applied, but they should not be regarded as biologically validated resilient or susceptible populations. Cross-method convergence measures robustness to analytical definition; it does not establish construct validity. This is also why method-specific terminology is important. Threshold-based stress responses, latent mixture classes, Two-Step partitions and Louvain communities are not equivalent statistical objects, even when they can be directionally aligned for comparison.

The same framework argues against validating resilience exclusively within the NTT. Behavioural stratification and biological validation represent distinct experimental stages. Bhatnagar emphasised that resilience is properly understood as a trajectory: an initial disruption followed by recovery towards previous functioning differs biologically from resistance, in which little disturbance occurs in the first place, even though both may culminate in similar endpoints (Bhatnagar, 2021). Cross-sectional stratification has been productive in preclinical work but cannot distinguish these trajectories and may therefore assign the same apparent outcome to biologically different processes (Bhatnagar, 2021). Our use of the term resilient-like is intended to preserve this distinction. The 22 reference-integrated fish identified here have a post-UCS behavioural organisation compatible with an operational resilient-like state; the design cannot determine whether they resisted behavioural perturbation, deviated and recovered, or achieved a compensated state through adaptation. Demonstrating resilience in the stronger temporal sense would require repeated measures capable of resolving baseline, perturbation and recovery.

Bhatnagar further noted that resilience in one domain can coexist with dysfunction in another and advocated multidimensional or multimodal measurement (Bhatnagar, 2021). Weavers similarly conceptualised biological resilience as a dynamic property spanning molecular, cellular, physiological and organismal levels and argued that mechanistic understanding requires integration across these scales (Weavers, 2024). Accordingly, the behavioural archetypes identified here should be regarded as experimentally testable hypotheses about organismal state. Their validation should include independent behavioural domains such as memory and social behaviour, alongside endocrine and physiological measures. Cortisol and HPI-axis dynamics are obvious candidates in the UCS model (Gallas-Lopes et al., 2023; Marcon et al., 2018a; Piato et al., 2011), but they should be used as independent outcomes rather than incorporated retrospectively into the behavioural definition. Measures involving glucocorticoid signalling (Piato et al., 2011), including GR and potentially MR/GR balance (Espigares et al., 2025), redox state (Marcon et al., 2018b, 2018a; Mocelin et al., 2019), inflammatory signalling (Marcon et al., 2016), neurotrophic and neuroplasticity-related processes, including BDNF signalling and adult neurogenesis (Chakravarty et al., 2013; Manuel et al., 2014; Song et al., 2018), and structural endpoints such as dendritic spine density or synaptic and neuronal morphology (Kumar et al., 2023; Pradhan et al., 2022; Song et al., 2018) may subsequently become valuable mechanistic readouts if supported by explicit hypotheses. Importantly, no single molecular marker should be expected necessarily to “normalise” in a resilient-like animal; maintenance of organismal function may itself depend on compensatory molecular adaptations.

This distinction between phenotypic identification and mechanistic explanation is supported by work in other experimental systems. Mueller et al. integrated locomotion, working memory, social preference and prepulse inhibition in a Two-Step analysis of offspring exposed to maternal immune activation, identifying exposed animals whose behavioural profiles resembled the reference control cluster and others showing overt multidomain impairment (Mueller et al., 2021). Behavioural stratification was subsequently used to examine transcriptional and neuroanatomical differences. Notably, animals classified as behaviourally resilient still showed CNS transcriptional alterations relative to non-exposed controls, which the authors interpreted as potentially consistent with allostatic adaptation rather than simple absence of biological effect (Mueller et al., 2021). In zebrafish larvae, Swaminathan et al. used the dynamics of recovery from stress to distinguish resilient from susceptible individuals and subsequently linked those phenotypes to distinct transcriptional responses and experimentally manipulable factors, including neuropeptide and complement pathways (Swaminathan et al., 2023). Related rodent studies have likewise shown that stress-exposed populations can segregate into different behavioural response phenotypes associated with distinct circuit, cellular or molecular adaptations (Anacker et al., 2018; Friedman et al., 2014; Krishnan et al., 2007). These studies differ substantially in species, stressor, behavioural criteria and definition of successful adaptation, and should not be interpreted as evidence for a universal molecular signature of resilience. Their common methodological principle is more relevant here: mechanistic comparisons become more interpretable when phenotypic heterogeneity is resolved before biological differences are sought.

This sequence has direct implications for the broader experimental programme. The present study addresses the phenotypic problem that precedes mechanistic analysis: how to identify, among genetically and experimentally comparable UCS-exposed zebrafish, individuals occupying reproducibly different regions of a complex behavioural response space without assuming in advance that a single endpoint or threshold constitutes resilience. Once such operational classifications are prospectively replicated and confronted with independent behavioural and physiological measures, individually traceable resilient- and susceptible-like fish can provide the substrate for targeted cellular hypotheses. In that setting, analyses of proteostasis and the unfolded protein response (Díaz-Hung and Hetz, 2022), glucocorticoid signalling (Spencer-Segal and Akil, 2019), redox regulation (Gao et al., 2025), inflammatory pathways (Dudek et al., 2021), neuronal morphology (Michaelson et al., 2020; Prévôt et al., 2017) or transcriptional programmes (Krishnan et al., 2007; Swaminathan et al., 2023; Willmore et al., 2022) become tests of mechanisms associated with a previously defined phenotype rather than *post hoc* criteria used to create that phenotype. The distinction is fundamental: the behavioural phenotype is not itself the cellular mechanism of resilience; it is the organism-level stratification required to ask mechanistic questions coherently. In this sense, behavioural classification provides the experimental bridge between heterogeneous responses to chronic stress and subsequent investigation of the neuroendocrine, cellular and molecular processes associated with successful or unsuccessful adaptation.

The present findings therefore suggest a broader interpretation of the NTT in resilience research. Novelty-evoked exploratory space use is neither a unitary measure of anxiety nor a direct assay of resilience. It is a context-dependent, temporally evolving and multivariate expression of how an individual deploys locomotion, space use and behavioural transitions after imposed novelty. UCS altered the distribution of individuals within this behavioural architecture, but the architecture itself remained largely overlapping with the unstressed reference when represented by the four variables available here. Within the exposed population, graph-based and alternative classifications could identify reproducible extremes, yet their biological meaning depended on the control reference and on the analytical definition applied. Future progress will therefore require richer behavioural sampling, preservation of temporal and spatial structure, explicit separation of resilience from resistance, and validation across independent biological domains. If those requirements are met, behavioural stratification can serve not as the endpoint of resilience research but as its experimental entry point: a means of identifying individuals in whom neuroendocrine, cellular, molecular and systems-level mechanisms associated with differential adaptation to chronic stress can subsequently be tested without using those same mechanisms retrospectively to define the phenotype.

## 5. Conclusion

Novelty-evoked exploratory space use in adult zebrafish was organised predominantly as a continuous multivariate behavioural architecture, within which UCS produced a significant redistribution towards lower exploration without evidence of a distinct behavioural space within the dimensions examined. *De novo* analysis of the UCS-exposed population revealed heterogeneous post-stress configurations, but higher exploratory expression alone was insufficient to define resilient-like individuals; its operational identification required integration with the unstressed reference architecture. Classification was further method-dependent, although a subset of zebrafish showed consistent cross-method assignments. These findings support reference-integrated, multivariate behavioural stratification as an operational approach for identifying resilient- and susceptible-like candidates for subsequent validation using independent physiological, cellular and molecular endpoints.

## Authors’ Contribution and Agreement

Conceptualization: F Fuentealba-Villarroel, AP Herrmann, D Zancan, and A Piato. Data curation: F Fuentealba-Villarroel and M Gallas-Lopes. Formal analysis: F Fuentealba-Villarroel, LM Bastos, and M Gallas-Lopes. Funding acquisition: F Fuentealba-Villarroel and A Piato.

Investigation: F Fuentealba-Villarroel and LM Bastos. Methodology: F Fuentealba-Villarroel, LM Bastos, and M Gallas-Lopes. Project administration: F Fuentealba-Villarroel and A Piato.

Resources: A Piato. Supervision: A Piato. Writing - original draft: F Fuentealba-Villarroel. Writing - review & editing: F Fuentealba-Villarroel, LM Bastos, M Gallas-Lopes, AP Herrmann, D Zancan, and A Piato.

The authors confirm that the work is original, has not been published previously, and is not under consideration for publication elsewhere. All necessary institutional permissions for the use and publication of the research data have been obtained, where applicable. All authors agree to be accountable for all aspects of the work and to ensure that any questions concerning its accuracy or integrity are appropriately investigated and resolved.

## Data availability

The original behavioural dataset, data dictionary, analysis scripts, computational environment documentation, SPSS Two-Step clustering materials and independent replication protocol are publicly available through Zenodo (https://doi.org/10.5281/zenodo.23425599). Processed datasets and analytical outputs can be regenerated using the documented computational workflows. The version-controlled source repository is available at GitHub: https://github.com/MisfoldERMind/zebrafish-ucs-exploratory-archetypes.

The broader research programme within which this retrospective analysis was conducted is publicly documented in the Open Science Framework (Fuentealba-Villarroel et al., 2025), which provides a time-stamped record of the overarching scientific rationale and analytical framework.

## References

1. Anacker, C., Luna, V.M., Stevens, G.S., Millette, A., Shores, R., Jimenez, J.C., Chen, B., Hen, R., 2018. Hippocampal neurogenesis confers stress resilience by inhibiting the ventral dentate gyrus. Nature 559, 98–102. 10.1038/s41586-018-0262-4

2. Anderson, M.J., 2017. Permutational Multivariate Analysis of Variance (PERMANOVA), in: Wiley StatsRef: Statistics Reference Online. John Wiley & Sons, Ltd, pp. 1–15. 10.1002/9781118445112.stat07841

3. Anderson, M.J., 2006. Distance-Based Tests for Homogeneity of Multivariate Dispersions. Biometrics 62, 245–253. 10.1111/j.1541-0420.2005.00440.x

4. Anderson, M.J., 2001. A new method for non-parametric multivariate analysis of variance. Austral Ecol. 26, 32–46. 10.1111/j.1442-9993.2001.01070.pp.x

5. Beigloo, F., Davidson, C.J., Gjonaj, J., Perrine, S.A., Kenney, J.W., 2024. Individual differences in the boldness of female zebrafish are associated with alterations in serotonin function. J. Exp. Biol. 227, jeb247483. 10.1242/jeb.247483

6. Benassi, M., Garofalo, S., Ambrosini, F., Sant’Angelo, R.P., Raggini, R., De Paoli, G., Ravani, C., Giovagnoli, S., Orsoni, M., Piraccini, G., 2020. Using Two-Step Cluster Analysis and Latent Class Cluster Analysis to Classify the Cognitive Heterogeneity of Cross-Diagnostic Psychiatric Inpatients. Front. Psychol. 11, 1085. 10.3389/fpsyg.2020.01085

7. Bertelli, P.R., Mocelin, R., Marcon, M., Sachett, A., Gomez, R., Rosa, A.R., Herrmann, A.P., Piato, A., 2021. Anti-stress effects of the glucagon-like peptide-1 receptor agonist liraglutide in zebrafish. Prog. Neuropsychopharmacol. Biol. Psychiatry 111, 110388. 10.1016/j.pnpbp.2021.110388

8. Bhatnagar, S., 2021. Rethinking stress resilience. Trends Neurosci. 44, 936–945. 10.1016/j.tins.2021.09.005

9. Blondel, V.D., Guillaume, J.-L., Lambiotte, R., Lefebvre, E., 2008. Fast unfolding of communities in large networks. J. Stat. Mech. Theory Exp. 2008, P10008. 10.1088/1742-5468/2008/10/P10008

10. Chakravarty, S., Reddy, B.R., Sudhakar, S.R., Saxena, S., Das, T., Meghah, V., Brahmendra Swamy, C.V., Kumar, A., Idris, M.M., 2013. Chronic Unpredictable Stress (CUS)-Induced Anxiety and Related Mood Disorders in a Zebrafish Model: Altered Brain Proteome Profile Implicates Mitochondrial Dysfunction. PLoS ONE 8, e63302. 10.1371/journal.pone.0063302

11. Csárdi, G., Nepusz, T., Traag, V., Horvát, S., Zanini, F., Noom, D., Müller, K., Antonov, M., Initiative, C.Z., Schoch, D., Salmon, M., details, R.C. igraph author, 2026. igraph: Network Analysis and Visualization. 10.5281/zenodo.7682609

12. Díaz-Hung, M.-L., Hetz, C., 2022. Proteostasis and resilience: on the interphase between individual’s and intracellular stress. Trends Endocrinol. Metab. TEM 33, 305–317. 10.1016/j.tem.2022.02.003

13. Dudek, K.A., Dion-Albert, L., Kaufmann, F.N., Tuck, E., Lebel, M., Menard, C., 2021. Neurobiology of resilience in depression: immune and vascular insights from human and animal studies. Eur. J. Neurosci. 53, 183–221. 10.1111/ejn.14547

14. Dunn, T.W., Mu, Y., Narayan, S., Randlett, O., Naumann, E.A., Yang, C.-T., Schier, A.F., Freeman, J., Engert, F., Ahrens, M.B., 2016. Brain-wide mapping of neural activity controlling zebrafish exploratory locomotion. eLife 5, e12741. 10.7554/eLife.12741

15. Espigares, F., Alvarado, M.V., Abad-Tortosa, D., Varela, S.A.M., Sobral, D., Faísca, P., Paixão, T., Oliveira, R.F., 2025. Optimistic and pessimistic cognitive judgement bias modulates the stress response and cancer progression in zebrafish. Transl. Psychiatry 15, 111. 10.1038/s41398-025-03311-9

16. Fernando, G.C.P., Khansari, A.R., Tort, L., 2024. Response to chronic crowding stress in shy and bold behavioral groups of male and female zebrafish. Am. J. Physiol.-Regul. Integr. Comp. Physiol. 327, R275–R290. 10.1152/ajpregu.00041.2024

17. Fontana, B.D., Moraes, H.S., Pretzel, C.W., Mohammed, K.A., Canzian, J., Bonan, C.D., Parker, M.O., Rosemberg, D.B., 2025. Individual Differences in Boldness Influence Working Memory and Stress-Induced Repetitive Behaviors in Zebrafish. Eur. J. Neurosci. 62, e70173. 10.1111/ejn.70173

18. Fraley, C., Raftery, A.E., 2007. Bayesian Regularization for Normal Mixture Estimation and Model-Based Clustering. J. Classif. 24, 155–181. 10.1007/s00357-007-0004-5

19. Friedman, A.K., Walsh, J.J., Juarez, B., Ku, S.M., Chaudhury, D., Wang, J., Li, X., Dietz, D.M., Pan, N., Vialou, V.F., Neve, R.L., Yue, Z., Han, M.-H., 2014. Enhancing depression mechanisms in midbrain dopamine neurons achieves homeostatic resilience. Science 344, 313–319. 10.1126/science.1249240

20. Fuentealba-Villarroel, F., Herrmann, A.P., Zancan, D., Piato, A., 2025. Targeting Protein Folding Pathways in Stress-Resilient Zebrafish: Insights into the Unfolded Protein Response - Experiment Preregistration [WWW Document]. Open Sci. Framew. 10.17605/OSF.IO/Z5NRM

21. Gallas-Lopes, M., Bastos, L.M., Benvenutti, R., Panzenhagen, A.C., Piato, A., Herrmann, A.P., 2023. Systematic review and meta-analysis of 10 years of unpredictable chronic stress in zebrafish. Lab Anim. 52, 229–246. 10.1038/s41684-023-01239-5

22. Gallas-Lopes, M., Herrmann, A.P., Piato, A., 2026. A Standardized Unpredictable Chronic Stress Protocol for Adult Zebrafish. Curr. Protoc. 6, e70376. 10.1002/cpz1.70376

23. Gamer, M., Lemon, J., Fellows, I., Singh, P., 2026. irr: Various Coefficients of Interrater Reliability and Agreement. 10.32614/CRAN.package.irr

24. Gao, L., Wang, B., Cui, X., Xia, L., Li, X., Figueredo, Y.N., Li, D., Liu, K., Wang, H., Jin, M., 2025. Neochlorogenic acid ameliorates Alzheimer’s disease-like pathology via scavenging oxidative stress and restoring blood-brain barrier function in zebrafish. Prog. Neuropsychopharmacol. Biol. Psychiatry 138, 111334. 10.1016/j.pnpbp.2025.111334

25. Gerlai, R., Fernandes, Y., Pereira, T., 2009. Zebrafish (*Danio rerio*) responds to the animated image of a predator: Towards the development of an automated aversive task. Behav. Brain Res. 201, 318–324. 10.1016/j.bbr.2009.03.003

26. Golushko, N.I., Matrynov, D., Galstyan, D.S., Apukhtin, K.V., de Abreu, M.S., Yang, L., Stewart, A.M., Kalueff, A.V., 2025. Understanding (and appreciating) behavioral complexity of zebrafish novel tank assays. Behav. Processes 230, 105230. 10.1016/j.beproc.2025.105230

27. Hubená, P., Benrejdal, L., Brodin, D., Axling, J., Sarma, O.S., Bergman, P., Winberg, S., 2026. Effects of Stress Coping Styles and Social Defeat on Zebrafish Behaviour and Brain Transcriptomics. Neurosci. Bull. 42, 989–1003. 10.1007/s12264-025-01506-0 IBM Corp., 2025. IBM SPSS Statistics for Windows.

28. Kassambara, A., 2026. rstatix: Pipe-Friendly Framework for Basic Statistical Tests. 10.32614/CRAN.package.rstatix

29. Kenney, J.W., Steadman, P.E., Young, O., Shi, M.T., Polanco, M., Dubaishi, S., Covert, K., Mueller, T., Frankland, P.W., 2021. A 3D adult zebrafish brain atlas (AZBA) for the digital age. eLife 10, e69988. 10.7554/eLife.69988

30. Krishnan, V., Han, M.-H., Graham, D.L., Berton, O., Renthal, W., Russo, S.J., LaPlant, Q., Graham, A., Lutter, M., Lagace, D.C., Ghose, S., Reister, R., Tannous, P., Green, T.A., Neve, R.L., Chakravarty, S., Kumar, A., Eisch, A.J., Self, D.W., Lee, F.S., Tamminga, C.A., Cooper, D.C., Gershenfeld, H.K., Nestler, E.J., 2007. Molecular Adaptations Underlying Susceptibility and Resistance to Social Defeat in Brain Reward Regions. Cell 131, 391–404. 10.1016/j.cell.2007.09.018

31. Kumar, H., Garg, V., Kaur, K., Kaur, R., 2023. Role of Spirulina in Structural Remodeling of Synapse in Telencephalon of Chronic Unpredictable Stress Model of Zebrafish. Ann. Neurosci. 30, 236–241. 10.1177/09727531231166202

32. Leong, N., Mujar, E.K., Mazlan, A.N., Chong, C.M., Christianus, A., Alresheedi, M.T., Ng, E.K., Yaacob, M.H., Mahdi, M.A., 2025. Discrimination of hypoxia tolerant and intolerant hybrid groupers using surface-enhanced Raman spectroscopy of epidermal mucus. Aquaculture 598, 742014. 10.1016/j.aquaculture.2024.742014

33. Maechler, M., Rousseeuw, P., Struyf, A., Hubert, M., Hornik, K., 2026. cluster: “Finding Groups in Data”: Cluster Analysis Extended Rousseeuw et al. 10.32614/CRAN.package.cluster

34. Manuel, R., Gorissen, M., Zethof, J., Ebbesson, L.O.E., van de Vis, H., Flik, G., van den Bos, R., 2014. Unpredictable chronic stress decreases inhibitory avoidance learning in Tuebingen long-fin zebrafish: stronger effects in the resting phase than in the active phase. J. Exp. Biol. 217, 3919–3928. 10.1242/jeb.109736

35. Marcon, M., Herrmann, A.P., Mocelin, R., Rambo, C.L., Koakoski, G., Abreu, M.S., Conterato, G.M.M., Kist, L.W., Bogo, M.R., Zanatta, L., Barcellos, L.J.G., Piato, A.L., 2016. Prevention of unpredictable chronic stress-related phenomena in zebrafish exposed to bromazepam, fluoxetine and nortriptyline. Psychopharmacology (Berl.) 233, 3815–3824. 10.1007/s00213-016-4408-5

36. Marcon, M., Mocelin, R., Benvenutti, R., Costa, T., Herrmann, A.P., de Oliveira, D.L., Koakoski, G., Barcellos, L.J.G., Piato, A., 2018a. Environmental enrichment modulates the response to chronic stress in zebrafish. J. Exp. Biol. 221, jeb176735. 10.1242/jeb.176735

37. Marcon, M., Mocelin, R., De Oliveira, D.L., Da Rosa Araujo, A.S., Herrmann, A.P., Piato, A., 2019. Acetyl-L-carnitine as a putative candidate for the treatment of stress-related psychiatric disorders: Novel evidence from a zebrafish model. Neuropharmacology 150, 145–152. 10.1016/j.neuropharm.2019.03.024

38. Marcon, M., Mocelin, R., Sachett, A., Siebel, A.M., Herrmann, A.P., Piato, A., 2018b. Enriched environment prevents oxidative stress in zebrafish submitted to unpredictable chronic stress. PeerJ 6, e5136. 10.7717/peerj.5136

39. Michaelson, S.D., Miranda Tapia, A.P., McKinty, A., Silveira Villarroel, H., Mackay, J.P., Urban, J.H., Colmers, W.F., 2020. Contribution of NPY Y5 Receptors to the Reversible Structural Remodeling of Basolateral Amygdala Dendrites in Male Rats Associated with NPY-Mediated Stress Resilience. J. Neurosci. 40, 3231–3249. 10.1523/JNEUROSCI.2621-19.2020

40. Mocelin, R., Marcon, M., D’ambros, S., Mattos, J., Sachett, A., Siebel, A.M., Herrmann, A.P., Piato, A., 2019. N-Acetylcysteine Reverses Anxiety and Oxidative Damage Induced by Unpredictable Chronic Stress in Zebrafish. Mol. Neurobiol. 56, 1188–1195. 10.1007/s12035-018-1165-y

41. Moretz, J.A., Martins, E.P., Robison, B.D., 2007. The effects of early and adult social environment on zebrafish (Danio rerio) behavior. Environ. Biol. Fishes 80, 91–101. 10.1007/s10641-006-9122-4

42. Mueller, F.S., Scarborough, J., Schalbetter, S.M., Richetto, J., Kim, E., Couch, A., Yee, Y., Lerch, J.P., Vernon, A.C., Weber-Stadlbauer, U., Meyer, U., 2021. Behavioral, neuroanatomical, and molecular correlates of resilience and susceptibility to maternal immune activation. Mol. Psychiatry 26, 396–410. 10.1038/s41380-020-00952-8

43. Müller, K., Wickham, H., Bryan, J., 2026. tibble: Simple Data Frames.

44. Neves, N.A., Gallas-Lopes, M., Patelli-Alves, A., Müller, D.V., Bastos, L.M., Stahlhofer-Buss, T., Herrmann, A.P., Piato, A., 2026. Unpredictable chronic stress fails to induce robust changes in zebrafish social behavior. Neurosci. Lett. 878, 138594. 10.1016/j.neulet.2026.138594

45. Oksanen, J., Simpson, G.L., Blanchet, F.G., Kindt, R., Legendre, P., Minchin, P.R., O’Hara, R.B., Solymos, P., Stevens, M.H.H., Szoecs, E., Wagner, H., Barbour, M., Bedward, M., Bengtsson, H., Bolker, B., Borcard, D., Borman, T., Carvalho, G., Chirico, M., Caceres, M.D., Durand, S., Evangelista, H.B.A., FitzJohn, R., Friendly, M., Furneaux, B., Hannigan, G., Heida, I., Hill, M.O., Lahti, L., Martino, C., McGlinn, D., Ouellette, M.-H., Cunha, E.R., Smith, T., Stier, A., Braak, C.J.F.T., Weedon, J., 2026. vegan: Community Ecology Package. 10.32614/CRAN.package.vegan

46. Pedersen, T.L., 2025. patchwork: The Composer of Plots. 10.32614/CRAN.package.patchwork

47. Piato, Â.L., Capiotti, K.M., Tamborski, A.R., Oses, J.P., Barcellos, L.J.G., Bogo, M.R., Lara, D.R., Vianna, M.R., Bonan, C.D., 2011. Unpredictable chronic stress model in zebrafish (*Danio rerio*): Behavioral and physiological responses. Prog. Neuropsychopharmacol. Biol. Psychiatry, The Neurobiology of Neurodegenerative Disorder: From Basic to Clinical Research 35, 561–567. 10.1016/j.pnpbp.2010.12.018

48. Pradhan, L.K., Sahoo, P.K., Chauhan, N.R., Das, S.K., 2022. Temporal exposure to chronic unpredictable stress induces precocious neurobehavioral deficits by distorting neuromorphology and glutathione biosynthesis in zebrafish brain. Behav. Brain Res. 418, 113672. 10.1016/j.bbr.2021.113672

49. Prévôt, T.D., Gastambide, F., Viollet, C., Henkous, N., Martel, G., Epelbaum, J., Béracochéa, D., Guillou, J.-L., 2017. Roles of Hippocampal Somatostatin Receptor Subtypes in Stress Response and Emotionality. Neuropsychopharmacol. Off. Publ. Am. Coll. Neuropsychopharmacol. 42, 1647–1656. 10.1038/npp.2016.281

50. R Core Team, 2026. R: A language and environment for statistical computing.

51. Rajput, N., Parikh, K., Kenney, J.W., 2022. Beyond bold versus shy: Zebrafish exploratory behavior falls into several behavioral clusters and is influenced by strain and sex. Biol. Open 11, bio059443. 10.1242/bio.059443

52. Rajput, N., Parikh, K., Squires, A., Fields, K.K., Wong, M., Kanani, D., Kenney, J.W., 2025. Whole-Brain Mapping in Adult Zebrafish and Identification of the Functional Brain Network Underlying the Novel Tank Test. eNeuro 12, ENEURO.0382-24.2025. 10.1523/ENEURO.0382-24.2025

53. Rosemberg, D.B., Rico, E.P., Mussulini, B.H.M., Piato, Â.L., Calcagnotto, M.E., Bonan, C.D., Dias, R.D., Blaser, R.E., Souza, D.O., Oliveira, D.L. de, 2011. Differences in Spatio-Temporal Behavior of Zebrafish in the Open Tank Paradigm after a Short-Period Confinement into Dark and Bright Environments. PLOS ONE 6, e19397. 10.1371/journal.pone.0019397

54. Sachett, A., Gallas-Lopes, M., Benvenutti, R., Marcon, M., Aguiar, G.P.S., Herrmann, A.P., Oliveira, J.V., Siebel, A.M., Piato, A., 2022. Curcumin micronization by supercritical fluid: In vitro and in vivo biological relevance. Ind. Crops Prod. 177, 114501. 10.1016/j.indcrop.2021.114501

55. Schauberger, P., Walker, A., 2025. openxlsx: Read, Write and Edit xlsx Files. 10.32614/CRAN.package.openxlsx

56. Scrucca, L., Fraley, C., Murphy, T.B., Raftery, A.E., 2023. Model-based clustering, classification, and density estimation using mclust in R, First edition. ed, Chapman & Hall/CRC the R series. Chapman & Hall/CRC, Boca Raton, FL. 10.1201/9781003277965

57. Simpson, G.L., 2026. permute: Functions for Generating Restricted Permutations of Data. 10.32614/CRAN.package.permute

58. Slowikowski, K., 2026. ggrepel: Automatically Position Non-Overlapping Text Labels with “ggplot2.”

59. Song, C., Liu, B.-P., Zhang, Y.-P., Peng, Z., Wang, J., Collier, A.D., Echevarria, D.J., Savelieva, K.V., Lawrence, R.F., Rex, C.S., Meshalkina, D.A., Kalueff, A.V., 2018. Modeling consequences of prolonged strong unpredictable stress in zebrafish: Complex effects on behavior and physiology. Prog. Neuropsychopharmacol. Biol. Psychiatry 81, 384–394. 10.1016/j.pnpbp.2017.08.021

60. Speedie, N., Gerlai, R., 2008. Alarm substance induced behavioral responses in zebrafish (Danio rerio). Behav. Brain Res. 188, 168–177. 10.1016/j.bbr.2007.10.031

61. Spence, R., Fatema, M.K., Reichard, M., Huq, K.A., Wahab, M.A., Ahmed, Z.F., Smith, C., 2006. The distribution and habitat preferences of the zebrafish in Bangladesh. J. Fish Biol. 69, 1435–1448. 10.1111/j.1095-8649.2006.01206.x

62. Spence, R., Gerlach, G., Lawrence, C., Smith, C., 2008. The behaviour and ecology of the zebrafish, Danio rerio. Biol. Rev. 83, 13–34. 10.1111/j.1469-185X.2007.00030.x

63. Spencer-Segal, J.L., Akil, H., 2019. Glucocorticoids and resilience. Horm. Behav. 111, 131–134. 10.1016/j.yhbeh.2018.11.005

64. Stewart, A., Cachat, J., Wong, K., Gaikwad, S., Gilder, T., DiLeo, J., Chang, K., Utterback, E., Kalueff, A.V., 2010. Homebase behavior of zebrafish in novelty-based paradigms. Behav. Processes 85, 198–203. 10.1016/j.beproc.2010.07.009

65. Swaminathan, A., Gliksberg, M., Anbalagan, S., Wigoda, N., Levkowitz, G., 2023. Stress resilience is established during development and is regulated by complement factors. Cell Rep. 42, 111973. 10.1016/j.celrep.2022.111973

66. Trotha, J.W. von, Vernier, P., Bally-Cuif, L., 2014. Emotions and motivated behavior converge on an amygdala-like structure in the zebrafish. Eur. J. Neurosci. 40, 3302–3315. 10.1111/ejn.12692

67. Videlier, M., Cornette, R., Bonneaud, C., Herrel, A., 2015. Sexual differences in exploration behavior in *Xenopus tropicalis*. J. Exp. Biol. 218, 1733–1739. 10.1242/jeb.120618

68. Weavers, H., 2024. Biological resilience in health and disease. Dis. Model. Mech. 17, dmm050799. 10.1242/dmm.050799

69. Wickham, H., Chang, W., Henry, L., Pedersen, T.L., Takahashi, K., Wilke, C., Woo, K., Yutani, H., Dunnington, D., Brand, T. van den, 2026a. ggplot2: Create Elegant Data Visualisations Using the Grammar of Graphics.

70. Wickham, H., François, R., Henry, L., Müller, K., Vaughan, D., 2026b. dplyr: A Grammar of Data Manipulation.

71. Wickham, H., Vaughan, D., Girlich, M., 2025. tidyr: Tidy Messy Data.

72. Willmore, L., Cameron, C., Yang, J., Witten, I.B., Falkner, A.L., 2022. Behavioural and dopaminergic signatures of resilience. Nature 611, 124–132. 10.1038/s41586-022-05328-2

73. Wong, K., Elegante, M., Bartels, B., Elkhayat, S., Tien, D., Roy, S., Goodspeed, J., Suciu, C., Tan, J., Grimes, C., Chung, A., Rosenberg, M., Gaikwad, S., Denmark, A., Jackson, A., Kadri, F., Chung, K.M., Stewart, A., Gilder, T., Beeson, E., Zapolsky, I., Wu, N., Cachat, J., Kalueff, A.V., 2010. Analyzing habituation responses to novelty in zebrafish (*Danio rerio*). Behav. Brain Res. 208, 450–457. 10.1016/j.bbr.2009.12.023

74. Wright, D., Rimmer, L.B., Pritchard, V.L., Krause, J., Butlin, R.K., 2003. Inter and intra-population variation in shoaling and boldness in the zebrafish (Danio rerio). Naturwissenschaften 90, 374–377. 10.1007/s00114-003-0443-2

75. Wullimann, M.F., Rupp, B., Reichert, H., 1996. The brain of the zebrafish Danio rerio: a neuroanatomical atlas, in: Wullimann, M.F., Rupp, B., Reichert, H. (Eds.), Neuroanatomy of the Zebrafish Brain: A Topological Atlas. Birkhäuser, Basel, pp. 19–87. 10.1007/978-3-0348-8979-7_5

76. Yata, K., Aoshima, M., 2020. Geometric consistency of principal component scores for high-dimensional mixture models and its application. Scand. J. Stat. 47, 899–921. 10.1111/sjos.12432

