## Supplementary Material for "Unpredictable chronic stress redistributes novelty-evoked exploratory space use and yields method-dependent identification of resilient-like archetypes in zebrafish"

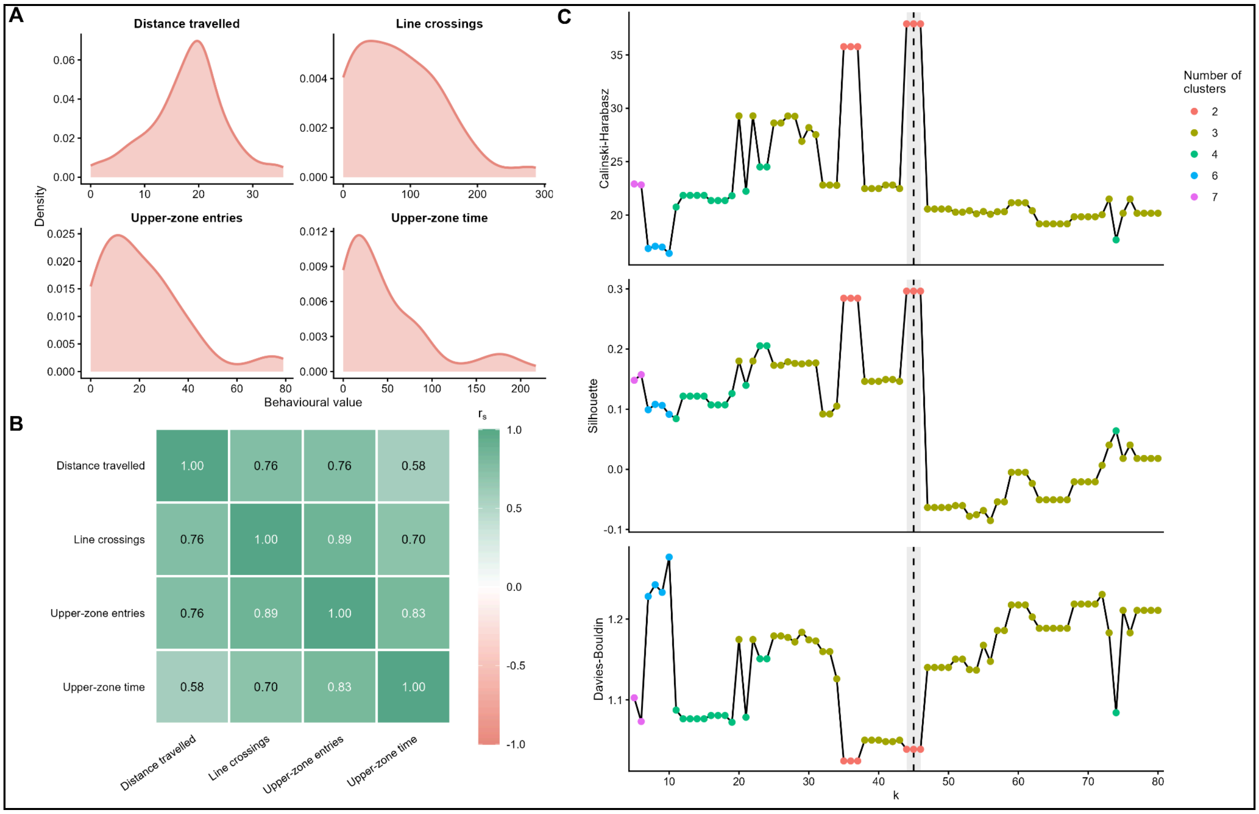


**Supplementary Figure S1. Behavioural distributions, covariation and neighbourhood-scale evaluation of Louvain partitions in the unstressed reference population of novelty-evoked exploratory space use.** (A) Kernel-density estimates of the four raw behavioural variables used to represent novelty-evoked exploratory space use in the 89 unstressed reference zebrafish: distance travelled, line crossings, upper-zone entries and upper-zone time. (B) Heatmap of pairwise Spearman correlations among the four behavioural variables. Cell colour and annotated values represent Spearman's rank correlation coefficient ($r_s$), with increasingly green tones indicating stronger positive associations. (C) Internal cluster-validation metrics obtained by varying the neighbourhood size from *k* = 5 to *k* = 80 in weighted, undirected union-*k*-nearest-neighbour graphs and applying Louvain community detection at each value of *k*. From top to bottom, the panels show the Calinski–Harabasz index, which quantifies between-cluster separation relative to within-cluster dispersion; mean silhouette width, which evaluates within-cluster cohesion relative to between-cluster separation; and the Davies–Bouldin index, for which lower values indicate better internal separation. Point colours indicate the number of clusters recovered at each neighbourhood size. The grey region denotes the exact-equivalence plateau from *k* = 44 to *k* = 46, within which identical two-cluster memberships were obtained, and the dashed vertical line marks *k* = 45, which was retained as the representative configuration for null-model and resampling analyses. The identical partition obtained at *k* = 46 was retained separately for subsequent projection analyses. The representative configuration was selected based on partition stability and complementary internal-validation metrics rather than the isolated optimum of any single index. These metrics were treated as descriptive optimisation criteria and not as inferential evidence of discrete behavioural populations.

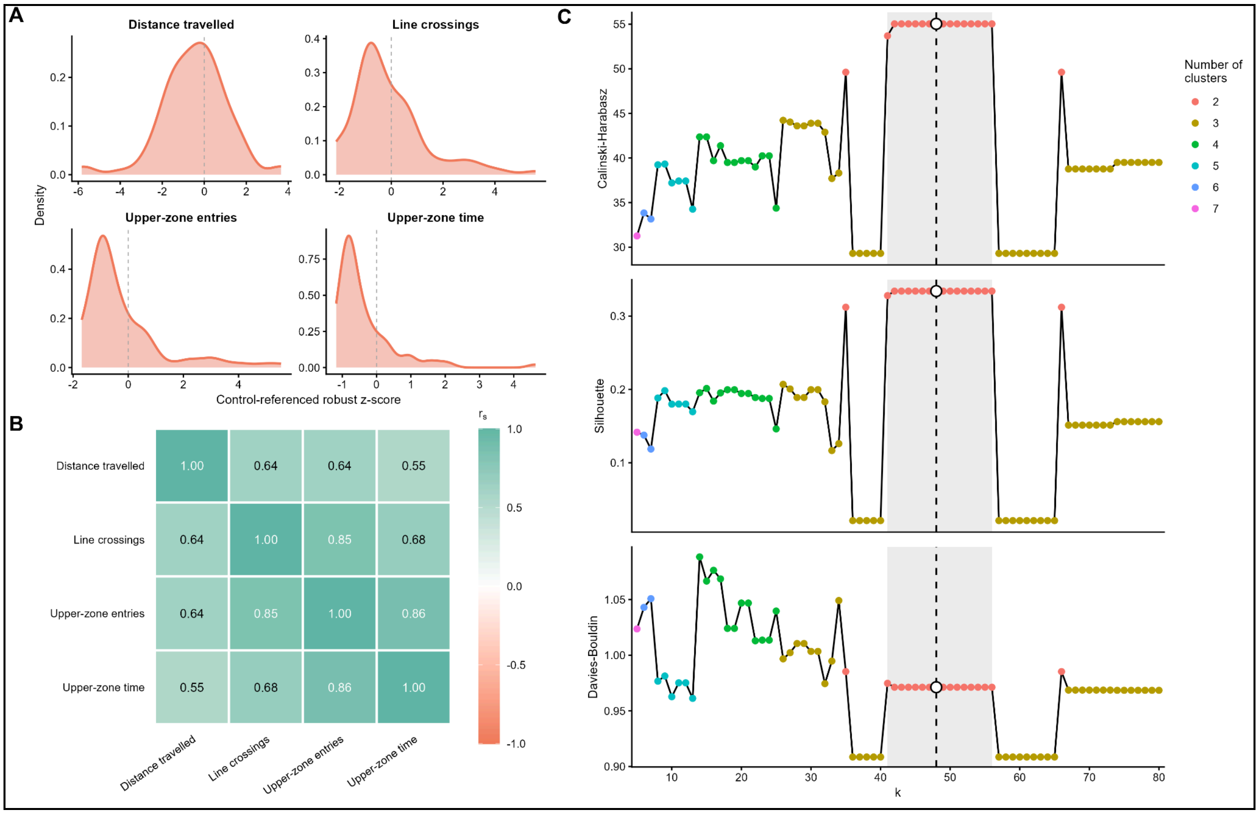


**Supplementary Figure S2. Behavioural distributions, covariation and neighbourhood-scale evaluation of *de novo* Louvain partitions of post-UCS novelty-evoked exploratory space use.** (A) Kernel-density estimates of the four control-referenced robust behavioural coordinates representing novelty-evoked exploratory space use in the UCS-exposed population (*n* = 89). Dashed vertical lines mark zero, corresponding to the cohort-specific control median on the inherited robust behavioural scale. (B) Spearman correlation matrix describing the dependence structure among the four behavioural variables within the UCS-exposed population. Cell colour and annotated values represent Spearman's rank correlation coefficient ($r_s$). Correlations ranged from $r_s$ = 0.55 for distance travelled and upper-zone time to $r_s$ = 0.86 for upper-zone entries and upper-zone time. (C) Internal cluster-validation metrics obtained by varying the neighbourhood size from *k* = 5 to *k* = 80 in weighted, undirected union-*k*-nearest-neighbour graphs and applying Louvain community detection at each value of *k*. From top to bottom, the panels show the Calinski–Harabasz index, mean silhouette width and Davies–Bouldin index. Point colours indicate the number of clusters recovered at each neighbourhood size. The grey-shaded region denotes the stable two-cluster plateau extending from *k* = 41 to *k* = 56, whereas the dashed vertical line indicates the representative solution selected at *k* = 48. At this solution, the Calinski–Harabasz index was 55.03, mean silhouette width was 0.334 and the Davies–Bouldin index was 0.971; open black circles identify the corresponding values. The representative configuration was selected based on stability across neighbouring partitions and complementary internal-validation metrics rather than the isolated optimum of any single index. These metrics were treated as descriptive optimisation criteria and not as inferential evidence of discrete post-UCS behavioural populations.

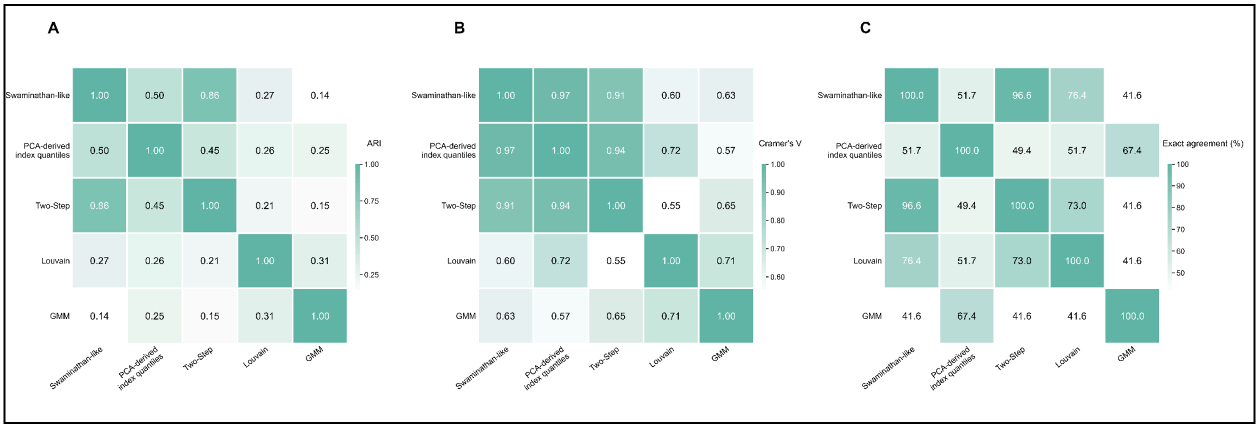


**Supplementary Figure S3. Pairwise agreement, partition similarity and association differ among analytical classification frameworks applied to post-UCS behavioural variation.** (A) Pairwise similarity between the classifications generated by the five analytical frameworks, quantified using the Adjusted Rand Index (ARI), which measures correspondence between individual-level assignments while accounting for agreement expected by chance. (B) Pairwise association between classifications, quantified using Cramér's *V*. (C) Exact individual-level agreement, expressed as the percentage of UCS-exposed zebrafish assigned to corresponding operational categories by each pair of frameworks. Across pairwise comparisons, exact agreement ranged from 41.6% to 96.6%, ARI ranged from 0.138 to 0.861 and Cramér's *V* ranged from 0.550 to 0.972. Swaminathan-like thresholding and Two-Step clustering showed the highest exact agreement and partition similarity (86/89 zebrafish; 96.6% exact agreement; ARI = 0.861; Cramér's $V = 0.915$; Fisher's exact test, $p = 1.55 \times 10^{-17}$). The lowest exact agreement occurred between GMM and both Swaminathan-like thresholding and Two-Step clustering (37/89 zebrafish; 41.6% in both comparisons), with corresponding ARI values of 0.138 and 0.153, respectively. Because the analytical frameworks differed in both category structure and operational definition, these pairwise statistics quantify concordance among classification schemes rather than equivalence of the underlying biological constructs. Diagonal values represent self-comparisons.
